# Empirical abundance distributions are more uneven than expected given their statistical baseline

**DOI:** 10.1101/2021.01.18.427126

**Authors:** Renata M. Diaz, Hao Ye, S. K. Morgan Ernest

## Abstract

Exploring and accounting for the emergent properties of ecosystems as complex systems is a promising horizon in the search for general processes to explain common ecological patterns. For example, the ubiquitous hollow-curve form of the species abundance distribution is frequently assumed to reflect ecological processes structuring communities, but can also emerge as a statistical phenomenon from the mathematical definition of an abundance distribution. Although the hollow curve may be a statistical artefact, ecological processes may induce subtle deviations between empirical species abundance distributions and their statistically most probable forms. These deviations may reflect biological processes operating on top of mathematical constraints and provide new avenues for advancing ecological theory. Examining ∼22,000 communities, we found that empirical SADs are highly uneven and dominated by rare species compared to their statistical baselines. Efforts to detect deviations may be less informative in small communities – those with few species or individuals – because these communities have poorly-resolved statistical baselines. The uneven nature of many empirical SADs demonstrates a path forward for leveraging complexity to understand ecological processes governing the distribution of abundance, while the issues posed by small communities illustrate the limitations of using this approach to study ecological patterns in small samples.

## Introduction

Ecological communities are complex systems made of numerous interacting entities subject to a vast array of processes operating in different contexts and at different scales (Levin 1992; Lawton 1999; Maurer 1999; Brown et al 2002; Nekola and Brown 2007; McGill 2019). One strategy for making sense of this inherent complexity is to identify patterns that occur consistently across many communities, and use these common phenomena to develop and test theories regarding general mechanisms that shape community structure (Brown and Maurer 1989; Maurer 1999; Lawton 1999; Gaston and Blackburn 2000; McGill 2019). Some of these patterns, however, can have counterintuitive emergent statistical properties (Frank 2009; 2019). Left unexamined, these properties can confound the interpretation of the observed patterns: what we interpret to be the result of generative mechanism may be an artifact of statistical constraints. However, when these properties are properly understood and accounted for, they can provide leverage for detecting and identifying the processes at work in a system (Jaynes 1957, Harte and Newman 2014).

The species abundance distribution (SAD) – the distribution of how all of the individuals in a community are divided among the species in that community – is a prime example of an ecological pattern that is both commonly invoked in the search for general processes, and subject to statistical constraints that have thus far complicated efforts to use it in this way (Nekola and Brown 2007; McGill et al. 2007; Locey and White 2013). The shape of the SAD is so consistent that it is often considered an ecological law (Preston 1948, 1962a, 1962b, 1980; Lawton 1999, McGill 2003, McGill et al. 2007). Across varied ecosystems and taxa, the species abundance distribution is dominated by a few very abundant species and a larger number of increasingly rare species, generating a distinctive hollow- or J-shaped curve when plotted with species rank on the x-axis and abundance on the y-axis (Fisher et al. 1943; McGill et al 2007). Community ecologists have used the SAD to test numerous theories regarding which biological processes are most important for structuring assemblages of species, by comparing theoretical predictions for the SAD to observed SADs (McGill 2003; McGill et al. 2007). However, this approach has proven inconclusive because many theories predict similar shapes for the SAD (McGill 2003; McGill et al. 2007), and even experimental manipulations generate little variation in the shape of the SAD (Supp and Ernest 2014). Investigating and accounting for the statistical considerations that constrain the shape of the SAD may open up new avenues for ecological interpretations of the SAD.

In fact, the nearly ubiquitous shape of the SAD may transcend ecological processes and instead reflect mathematical properties inherent to abundance distributions. Complex systems across domains ranging from economics to information technology often exhibit empirical abundance distributions with hollow-curve forms similar to ecological SADs (Shockley 1957; Gaston et al. 1993; Nekola and Brown 2007, Blonder et al. 2014; Keil et al. 2018). This suggests that the hollow curve is a common feature of abundance distributions and not necessarily an ecological phenomenon. Because the hollow-curve is observed in diverse systems and many theoretical generative processes converge to power-law or log-series abundance distributions (i.e. hollow curves) (Preston 1950; McGill 2003; Nekola and Brown 2007; Frank 2009; Frank 2019), approaches from statistical mechanics and complexity science may best explain the expected emergent shape for the distribution (Preston 1950; McGill 2003; Nekola and Brown 2007; Dewar and Porté 2008). Indeed, frameworks grounded in both entropy maximization (e.g. the Maximum Entropy Theory of Ecology; Harte et al. 2008, Harte 2011) and combinatorics (i.e. ‘the feasible set’; Locey and White 2013) generate realistic hollow curves via the random division of the total number of individuals in a community, *N*, into the total number of species present *S*. If the SAD is statistically inclined to be a hollow curve, the hollow-curve in itself may be of limited use for developing and testing ecological theories.

While SADs may be statistically constrained, this does not necessarily mean that they cannot be biologically informative. Biological factors may introduce subtle, but meaningful, deviations between observed SADs and the shapes of the SADs expected due to the mathematical constraints imposed by *S* and *N*, which we hereafter refer to as the “statistical baseline” (Locey and White 2013, Harte and Newman 2014). If the vast majority of mathematically achievable SADs for a community share a similar shape, an empirically observed SAD that deviates even slightly from this statistical baseline is unlikely to have emerged at random (Locey and White 2013), and may be the signature of a non-random – i.e., biological – process operating on the relative abundances of species (Harte and Newman 2014). If, over many communities, there are consistent deviations between observed SADs and their statistical baselines, these deviations can help evaluate and refine ecological theories. For example, the high prevalence of rare species in ecological communities has attracted considerable empirical and theoretical attention (e.g. Nee et al. 1991; Magurran and Henderson 2003), but it is unclear to what extent this phenomenon may derive from mathematical constraints on the SAD rather than ecological processes. If the prevalence of rare species in observed distributions consistently exceeds what would be expected to emerge from the statistical baseline, we would be prompted to look for ecological mechanisms promoting rarity. Candidate theories could then be evaluated based on how well their predictions for the rare tail of the SAD matched observed distributions. Thus, the *deviations* from the statistical baseline may enable us to detect strong ecological processes or evaluate theories (Harte and Newman 2014, Xiao et al. 2016).

Successfully interpreting SADs in this fashion depends on our capacity to detect and quantify deviations between empirical observations and statistical baselines, which requires metrics and computational approaches that allow us to quantify and interpret whatever deviations may exist. Here, we build upon the combinatoric approach developed by Locey and White (2013) to define and explore statistical baselines for SADs. For a given *N* (total number of individuals) and *S* (total number of species), there exists a finite set of possible distributions of individuals into species. Collectively, this set of possible SADs is the *feasible set*, with each possible SAD constituting a single element of the set. If an observed SAD is drawn at random from the feasible set, it is likely to have a shape similar to the shapes most common in the feasible set. The feasible set therefore allows us to define statistical baselines for assessing deviations between observed SADs and what is likely to occur due to mathematical constraints (Locey and White 2013).

The feasible set can also be used to explore how the characteristics of the statistical baseline, and the presence and nature of any deviations that occur, vary over ranges of values for S and N. Although most feasible sets are dominated by the hollow-curve shape, variation in S, N, and the ratio of N to S modulate the detailed attributes of the SADs in a feasible set (Locey and White 2013). For example, if the ratio of N to S is close to 1, all possible SADs are mathematically constrained to be fairly even (Locey and White 2013). Although an SAD that is very even would be highly unusual in most cases, it would be expected in this situation. The feasible set therefore allows us to appropriately calibrate our expectations for what types of observations would be surprising for an SAD given the specific constraints imposed by its S and N. Additionally, accounting for variation in the specificity, or vagueness, of the expectations derived from the statistical baseline may be critically important for disentangling the aspects of the SAD that can be attributed to statistical constraints from those that result from other processes. If the vast majority of mathematically possible SADs are similar in shape – generating a very specific, narrowly defined statistical baseline – then even small deviations between an observed SAD and this baseline can signal the operation of ecological processes. However, if many different shapes occur with more even frequency in the feasible set, the statistical baseline is less specific and less well defined, and our sensitivity for distinguishing biological signal from statistical constraints is greatly reduced. This is more likely to occur when the size of the community, in terms of *S* and *N*, is small, because in such cases the feasible set may be too small for a particular shape to emerge as the most common shape. These statistical baselines with broad distributions may therefore impede our ability to assess whether observed deviations are ecologically generated or expected to emerge randomly (Jaynes 1957). This general concern has been acknowledged in efforts to compare ecological observations to statistical baselines (Harte 2011, White et al. 2012, Locey and White 2013) but there has not yet been a quantification of these effects for the SAD or an identification of the range of community sizes most strongly affected. Because ecologists study the SAD for communities varying in size from the very small – *S* and *N* < 5 – to the enormous – *S* and *N* >> 1000 – identifying the community sizes for which we can and cannot confidently detect deviations from the statistical baseline is necessary to appropriately contextualize our interpretations.

Here we use the feasible set to define statistical baselines for empirical SADs for 22,000 communities of birds, mammals, trees, and miscellaneous other taxa. We then compare *observed* SADs to their corresponding statistical baselines and evaluate 1) if the shapes of observed SADs consistently deviate from their statistical baseline, 2) how the characteristics and specificity of the statistical baseline vary over ranges of *S* and *N*, and 3) whether this variation appears to be associated with variation in our capacity to detect deviations between observations and the corresponding baselines.

## Methods

Data and code for all of our analyses can be accessed at www.github.com/diazrenata/scadsanalysis.

### Datasets

We used a compilation of community abundance data for trees, birds, mammals, and miscellaneous additional taxa (White et al. 2012, Baldridge 2015, Baldridge 2016, data from Baldridge 2016). This compilation consists of cleaned and summarized community abundance data for trees obtained from the Forest Inventory and Analysis (Woudenberg et al 2010) and Gentry transects (Phillipes and Miller 2002), birds from the North American Breeding Bird Survey (Sauer et al. 2013), mammals from the Mammal Community Abundance Database (Thibault et al. 2011), and a variety of less commonly sampled taxa from the Miscellaneous Abundance Database (Baldridge 2015). Because characterizing the random expectation of the SAD is computationally intractable for very large communities, we filtered our datasets to remove 4 communities that had more than 40714 individuals, which was the largest community we successfully analyzed. We further filtered the FIA database. Of the 103,343 communities in FIA, 92,988 have fewer than 10 species. Rather than analyze all these small communities, we randomly selected 10,000 small communities to include in the analysis. We also included all FIA communities with more than 10 species, which added 10,355 FIA communities to the analysis and resulted in a total of 20,355 FIA communities. Finally, for sites that had repeated sampling over time, we followed White et al. (2012) and Baldridge (2016) and analyzed only a single, randomly selected, year of data, because samples taken from a single community at different time points are likely to covary. It should be noted that our analyses include data from the Mammal Community Database and Miscellaneous Abundance Database that were collected over longer timescales and cannot be disaggregated into finer units of time. Our final dataset consisted of ∼22,000 communities with S and N ranging from 2 to 250 and 4 to 40714, respectively (Figure 1). Details and code for the filtering process can be found in Appendix S1 in Supporting Information.

**Figure 1.**
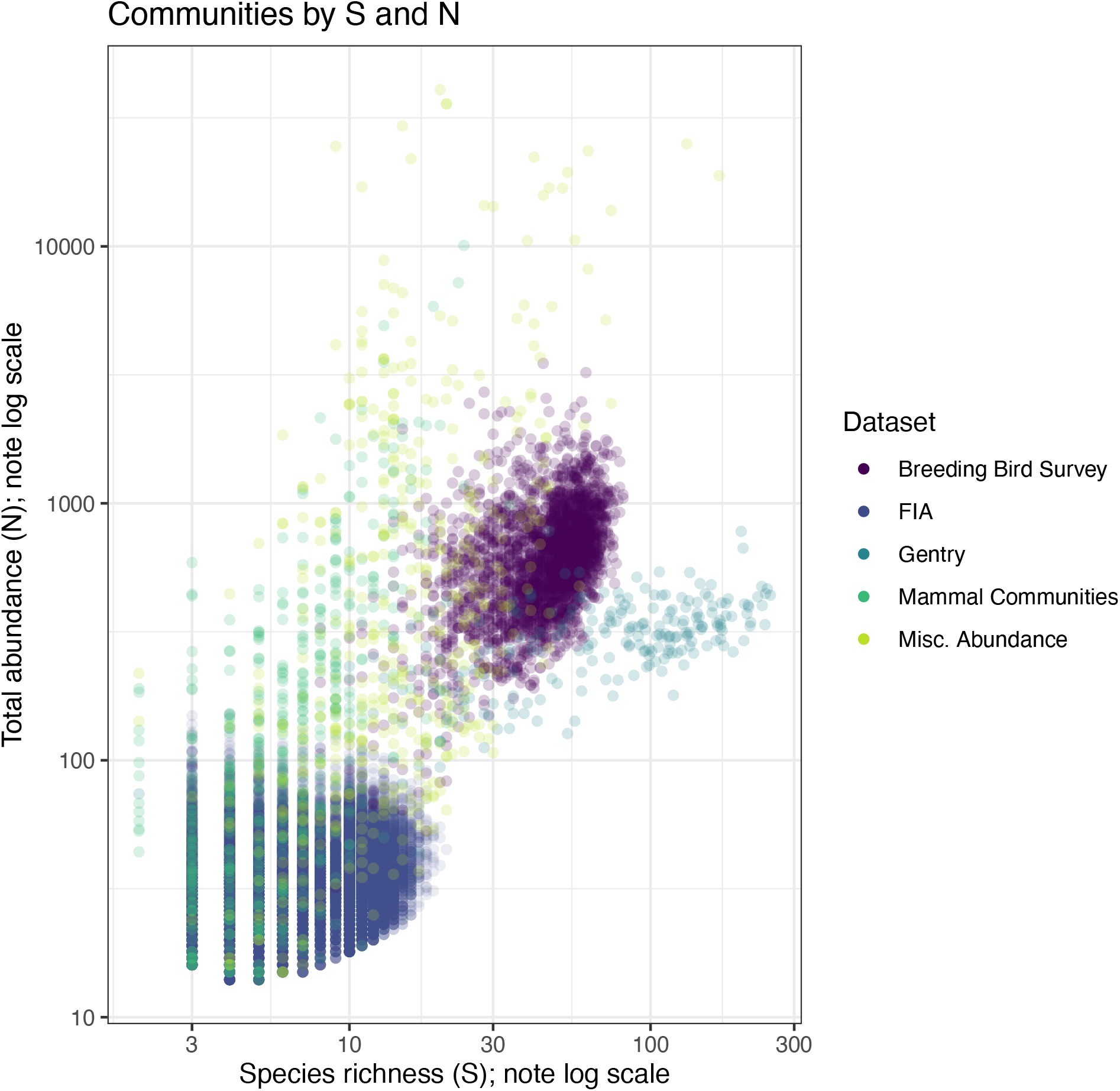
Distribution of communities from each dataset in terms of total abundance (N) and species richness (S). Communities range from few species and individuals (lower left corner) to speciose communities with many individuals (upper right). However, datasets are not evenly distributed across this state space due to differences in their sampling intensity, design, and underlying biology (e.g. productivity, regional richness, taxonomic group, or other factors that influence the capacity of a community to support large numbers of species and individuals). In particular, note that the FIA dataset comprises very small communities, and communities from the Gentry dataset are extreme in both their high species richness and low average abundance.

### Accounting for empirical sampling error

Because it is logistically impossible to exhaustively census all individuals present in most empirical systems, SADs derived from field sampling will inevitably be subject to some degree of sampling error (Bonar et al. 2011). Therefore, in addition to analyzing the raw SADs in our database, we employed two resampling schemes to test if, and how, different forms of observation error affect our results.

First, we explored the possibility that empirical sampling systematically undercounts the true number of rare species in a community (Preston 1948; Gotelli and Colwell 2011). Rare species are more likely to escape detection during sampling, leading to an underestimate of both the total species richness of a community and the proportion of species in the rare tail of the SAD (Preston 1948). We used a procedure based on species richness estimators to adjust for this possibility (see also Ulrich et al. 2010 for the use of richness estimators to distinguish between completely and incompletely censused communities). We computed the estimated richness for each community using the bias-corrected Chao and the ACE estimators (as implemented in the R package “vegan”; O’Hara 2005; Chiu et al 2014; Oksanen et al. 2020). To each of these richness estimates, we added one standard deviation of the estimate, and then took the mean of the two results. This yields a generous estimate of the true number of species in the system. If this estimate exceeded the observed species richness, we added the missing species each with abundance 1. These adjusted SADs allowed us to explore the consequences of undersampling rare species while making the smallest possible changes to S and N.

Second, we tested the sensitivity of our results to sampling variability across all species in the SAD – not just rare species - using subsampling. For each observed community, we constructed subsamples by randomly drawing 60% of the observed number of individuals from the total pool of individuals in the community, without regard to species and without replacement. The precise proportion of individuals drawn in each subsample should not dramatically affect the qualitative outcome. We selected 60% so as to introduce appreciable room for sampling error between the raw and subsampled SADs, but to produce subsampled SADs with N (and presumably S) in a comparable size range to the raw ones. Extremely small subsamples (e.g. 10%) could introduce complications related to small N and S that could obscure the effects of sampling error, while very large subsamples (e.g. 90%) could recapture the raw distributions too closely to be informative. We generated 10 resampled communities for each observed community.

We ran our computational pipeline using all raw SADs and all SADs adjusted for undersampling of rare species. Because the subsampling approach increased computational effort approximately tenfold, we analyzed all subsampled communities for the Mammal Community, Miscellaneous Abundance, and Gentry databases, but only a random subset of 300 (of 2773) communities from the Breeding Bird Survey and 2000 (of 20179) from the FIA – 1,000 with S < 10, and 1,000 with S >= 10.

### Generating the statistical baseline

We use the concept of the “feasible set” to establish a statistical baseline for the SAD (Locey and White 2013). For a given number of individuals *N*, there are a finite number of unique ways to partition those individuals into *S* species. The complete set of these unique partitions is the feasible set. In this approach, neither species nor individuals are distinguishable from each other; thus partitions are unique if and only if they differ in the number of species that have a particular abundance (Locey and White, 2013). Operationally, this means that for *S = 3* and *N = 9*, the SADs *(1, 3, 5)* and *(2, 2, 5)* count as distinct partitions, but *(1, 3, 5)* and *(3, 1, 5)* do not, because they each contain one species with an abundance 1, 3, and 5, respectively, and differ only in the *order* of the numbers. In the absence of justification for additional assumptions regarding the distinguishability of species and/or individuals, we adopted this simple set of assumptions that has previously been shown to generate realistic statistical baselines (Locey and White 2013).

While it is possible to list all possible partitions in the feasible set for small *S* and *N*, the size of the feasible set increases rapidly with *S* and *N*. An exhaustive characterization of the statistical properties of the feasible set for large *S* and *N* quickly becomes computationally intractable. This renders it necessary to draw samples from the feasible set, rather than enumerating all of its elements. Previous efforts in this vein (Locey and White 2013) have been constrained by the problem of unbiased sampling of large feasible sets. We developed an algorithm to efficiently and uniformly sample feasible sets even for large values of *S* and *N*. In brief, the algorithm takes a generative approach to sample the feasible set for a given combination of *S* and *N*, based on recurrence relations used to calculate the size of the feasible set. Let f(S, N) be the number of possible partitions of N individuals into exactly S species, i.e. the size of the feasible set for given values of S and N. Computation of f(S, N) can be achieved without enumerating the entire feasible set through the recurrence relation f(S, N) = f(S-1, N-1) + f(S, N-S) (originally documented in a 1742 letter from Euler to Bernoulli; 1862). For example, consider the feasible set with S = 3 and N = 7. For all possible partitions, either (a) at least one species has an abundance equal to 1, or (b) all of the species have abundance greater than 1. In the case of (a), removing one species with abundance equal to 1 must result in a partition of 6 individuals into 2 species. In fact, all of the unique partitions in (a) must have a corresponding unique partition in the feasible set for S = 2 and N = 6, and vice versa. In the case of (b), removing 1 individual from each species must result in a partition from the feasible set with S = 3 and N = 4. Here, all the partitions in (b) must have a corresponding unique partition in the feasible set with S = 3 and N = 4, and vice-versa. Therefore, f(3,7) = f(2,6) + f(3,4). By storing the values in a lookup table, f(S, N) can be calculated for increasing values of S and N through straightforward summation.

This recurrence relation also makes it possible to draw random samples from the feasible set without enumerating all possible partitions of N into S. For the example of S = 3 and N = 7, there are a total of 4 possible partitions (i.e. f(S, N) = 4). Because f(2, 6) = 3 and f(3, 4) = 1, we know that (a) 3 of the 4 partitions must correspond to a partition of the feasible set with S = 2 and N = 6 (but with a species of abundance equal to 1 removed), and (b) 1 of the 4 partitions must correspond to a partition of the feasible set with S = 3 and N = 4 (but with 1 individual removed from each species). Thus, we can determine the probability that a partition drawn at random from the feasible set for S = 3 and N = 4 is in case (a) – probability ¾ - or case (b) – probability ¼. To generate a partition in case (a), we sample a partition for S = 2 and N = 6 and then add a species with abundance equal to 1; for case (b), we sample a partition for S = 3 and N = 4 and then add 1 individual to each species. In this way, we use the recurrence relation to transform the problem of sampling from a large feasible set into the problem of sampling from a smaller, different feasible set. This procedure continues until a partition is uniquely determined, after which some back-transformation yields a unique partition for the feasible set of interest. A detailed description of the algorithm we use, based on a slightly different recurrence relation, is available in Appendix S2 and is implemented in the R package `feasiblesads` available on GitHub at www.github.com/diazrenata/feasiblesads.

For every community in our database, we drew 4000 samples from the feasible set to characterize the distribution of statistically probable shapes for the SAD. We filtered the 4000 samples to unique elements. For small values of S and N, it can be impossible or highly improbable for the 4000 samples from the feasible set to all be unique, but for large communities, all 4000 are usually unique. We refer to this as the sampled feasible set.

### Comparing observed SADs to their statistical baselines

We compared SADs to their statistical baselines using several metrics, including a general measure of dissimilarity, as well as skewness, Simpson’s evenness, Shannon’s index, and the proportion of rare species (species with abundance = 1). These metrics represent just a few of the vast array of possible summary metrics to describe the shape of the SAD, each of which emphasize different aspects of the distribution. In this first effort to compare empirical distributions to a statistical baseline, we selected a suite of complementary metrics and explored whether our overall results were consistent between metrics. By calculating these metrics for each the community’s sampled feasible set (see *Generating the statistical baseline*, above), we generated a portfolio of measures describing the shapes expected from randomly sampled SADs.

First, as a general characterization of whether observed SADs have rare or common shapes relative to their feasible sets, we computed a dissimilarity score comparing SADs to the central tendencies of their feasible sets (following Locey and White, 2013). We defined the degree of dissimilarity between two SADs with the same S and N as the proportion of individuals allocated to species with different abundances between the two SADs, calculated as:

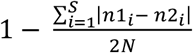

where *n1*_*i*_ is the abundance at rank *i* for one SAD and *n2*_*i*_ is the abundance at rank *i* for the other SAD. This value ranges from 0 to 1, with 1 being high dissimilarity. To find the central tendency of a given sampled feasible set, we identified the sampled SAD with the lowest mean dissimilarity compared to the rest of the SADs in the feasible set. We calculated the dissimilarity between every sample drawn from the feasible set and a random set of 500 other samples, using a subset of samples for comparisons because it is computationally impractical to make all pairwise comparisons between large numbers of samples. To assess whether an observed SAD was highly dissimilar to its central tendency, we calculated the degree of dissimilarity between the central tendency of the corresponding feasible set and all other samples from that feasible set, and between the central tendency and the observed SAD. Although the dissimilarity score is scaled from 0 to 1, the distributions of dissimilarity scores for samples from the feasible set can vary over broad ranges in S and N. We therefore used the percentile rank of the observed dissimilarity scores, relative to the distribution of dissimilarity scores from the corresponding sampled feasible sets, to quantify how likely or unlikely observed dissimilarity scores are across the range of S and N in our datasets. For a single community, an observed percentile score of 95 indicates that there is a 5% chance of drawing a value greater than the observed value from the distribution of values from the sampled feasible set. Aggregating across communities, if observed SADs reflect random draws from their feasible sets, their percentile rank values should be uniformly distributed from 0 to 100. However, if observed SADs are consistently more dissimilar to their feasible sets that expected at random, the percentile values will be disproportionately concentrated at high values. We used a one-tailed 95 confidence interval and tested whether the percentile values for the dissimilarity scores of observed SADs fell above 95 more than 5% of the time. We note that it is impossible for an observation fall above the 95^th^ percentile if there are fewer than 20 values in the sampled distribution. We therefore excluded from this analysis communities with fewer than 20 unique SADs in their feasible sets, yielding a total of 22,490 communities. Finally, note that, if the observed dissimilarity scores for individual communities are not systematically higher than the distributions of dissimilarity scores from the corresponding feasible sets, increasing the number of *communities* in the analysis will not increase the frequency of extreme percentile scores.

While the degree of dissimilarity between SADs and the central tendency of the feasible set provides an overall sense of deviations among possible SADs, it does not describe *how* observed SADs may differ from their feasible set. We therefore used a set of more targeted, ecologically interpretable metrics to explore how observed SADs compare to their feasible sets in their shape and proportion of rare species. We examined three metrics for the shape of the SAD - skewness, Simpson’s evenness (1-D), and Shannon’s index. Skewness measures the asymmetry of a distribution around its mean. The Simpson and Shannon indices are commonly used metrics for assessing how equitably abundance is distributed across species (Maurer and McGill 2011). We also calculated the proportion of rare species (species with abundance = 1) in each SAD, because the proportion of rare species in a community is comparable across different community sizes and is of special interest to ecologists.

As with the degree of dissimilarity score, to assess whether the shape of an observed SAD was statistically unlikely, we used percentile ranks to compare the observed values of the summary metrics to the distributions of values for those metrics obtained from each community’s sampled feasible set. The actual ranges and values of summary metrics vary widely over the ranges of S and N in our data and thus cannot directly compared, but percentile ranks are comparable across different community sizes and allow assessment across our entire dataset. We used two-tailed 95% intervals to test whether observed communities’ percentile values for each metric were disproportionately concentrated below 2.5 or above 97.5. In all cases, in testing for unusually high percentile scores, we defined the percentile score as the proportion of values in the sampled distribution strictly less than the observed value, while in testing for low values, we defined it as the proportion of sampled values less than or equal to the observed value. This ensured a conservative estimate of how extreme the observed values were relative to the sampled distribution. Because it is impossible for an observed percentile score to be above or below the 97.5^th^ or 2.5^th^ percentile if there are fewer than 40 values in the sample distribution, we excluded from these analyses communities with fewer than 40 SADs in their feasible sets. Finally, note that skewness, as implemented in the R package “e1071” (Meyer et al. 2019), always evaluates to 0 for distributions with only two species, and we therefore excluded those cases from analyses of skewness. Our final analysis included 21,395 communities for skewness and 21,403 communities for all other shape metrics.

### The narrowness of the expectation

We also used the distributions of dissimilarity scores and shape metrics to quantify the relative specificity of the statistical baseline, in order to assess when there could be challenges in determining whether observed communities differ from their statistical baselines. For an overall sense of how tightly elements of the feasible set were clustered around its central tendency, we calculated the mean dissimilarity score between all samples from a feasible set and the central tendency of that feasible set. For the shape metrics, we calculated a breadth index defined as the ratio of the range of values encompassed within a two-sided 95% density interval relative to the full range of values in the distribution (Figure 2). This breadth index for the statistical baseline ranges from 0 (a very narrow distribution and well-resolved baseline) to 1 (a very broad distribution), and is comparable across feasible sets for varying combinations of *S* and *N*. These approaches correspond qualitatively to more computationally-intensive approaches to measuring the self-similarity of the elements of feasible sets (see Appendix S3). We explored how the narrowness of the statistical baseline varies with the size of the feasible set and the ratio of N to S.

**Figure 2.**
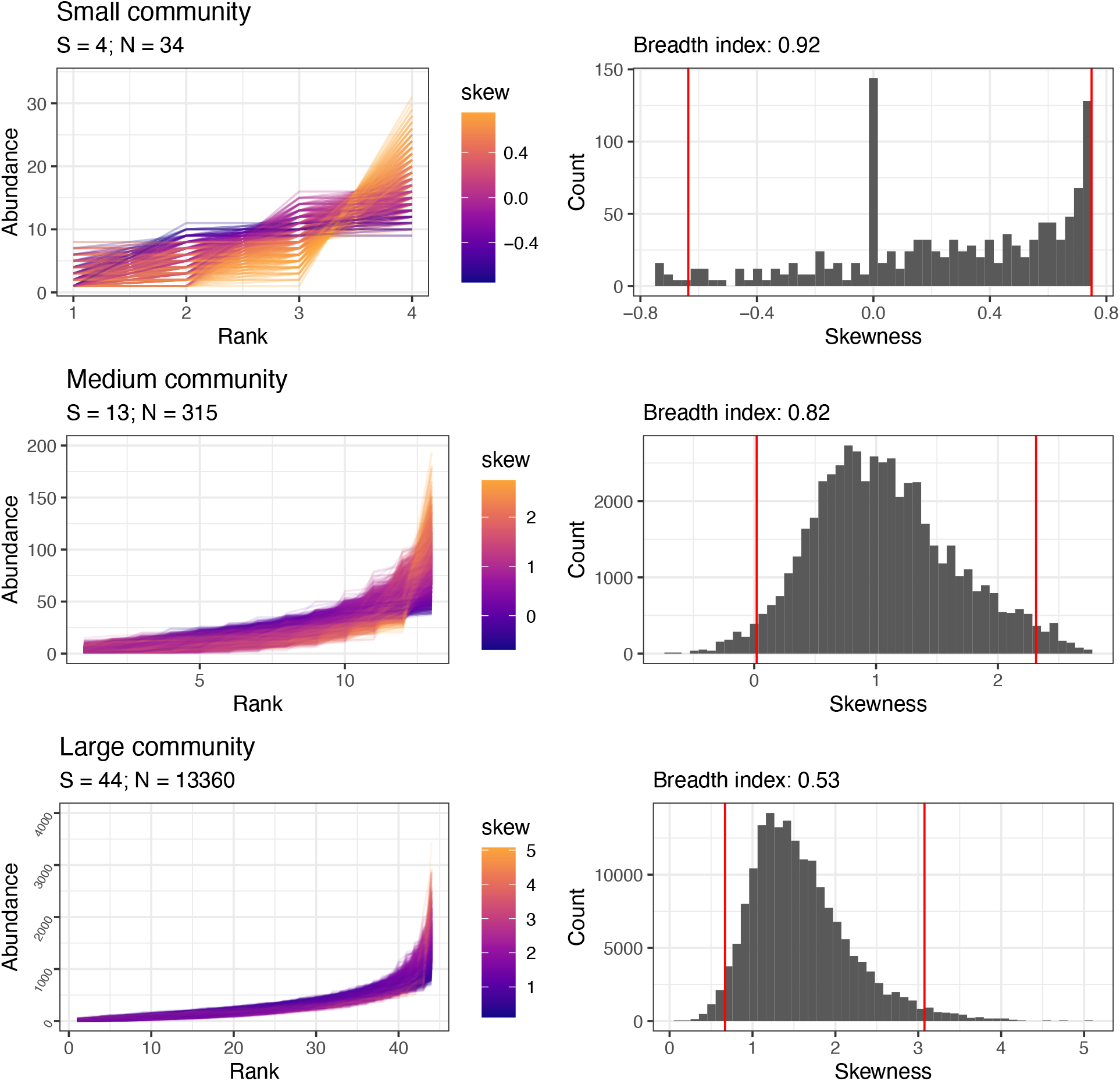
Large feasible sets may allow better detection of deviations from the statistical baseline by generating more specific, narrowly-defined baselines. We illustrate this phenomenon using 3 hypothetical communities: a small community (*S* = 4, *N* = 34; top row), an intermediate community (*S* = 13, *N* = 315; middle row), and a large community (*S*= 44, *N* = 13360; bottom row). The large communiity has approximately 6.59e+70 possible SADs in its feasible set, while the intermediate community has 1.001e+12 and the small community has only 297. For every SAD sampled from the feasible set (left column), we calculate the skewness (color scale) or other summary metrics (not shown). The distributions of these values (right column) constitute the statistical baseline. We define a “breadth index” as the ratio of the range encompassed in the two-tailed 95% density interval (distance between red lines, right), compared to the full range of values for the statistic (distance between the maximum and minimum values). As *S* and *N* increase, the size of the feasible set increases, resulting in a narrower statistical baseline (smaller breadth index) – thus enabling easier detection of deviations that may be the result of ecological processes affecting the SAD.

## Results

### Comparing observed SADs to their statistical baselines

For four of the five datasets we analyzed – BBS, Gentry, Mammal Communities, and Misc. Abund – observed SADs are more dissimilar to their statistical baselines than would be expected by chance (Figure 3). Combined over these four datasets, 29% of observed SADs are more dissimilar to the central tendency than are 95% of samples from the corresponding feasible sets (Table 1). If observed SADs reflected random draws from the feasible set, we would expect only 5% to be that dissimilar. These highly unlikely SADs have dissimilarity scores from 1.5 to 9.7 times greater than the mean dissimilarity between the central tendency and samples from the feasible set, an absolute increase ranging from .04 to .6 on a scale from 0-1 (Figure S4). These datasets also contain highly unlikely observed SADs in terms of their shape metrics. At random, roughly 2.5% of observed percentile scores for these metrics should be very high (>97.5) or very low (<2.5). Compared to their feasible sets, these four datasets contain a disproportionate number of communities with very low values for Simpson’s evenness and Shannon diversity, and very high skewness, relative to their feasible sets (Table 1). The Mammal Community and Miscellaneous Abundance databases also have high proportions of rare species, but this tendency is weaker for BBS and nonexistent for Gentry – in fact, the Gentry dataset has a high representation of sites with *low* proportions of rare species (20% of sites; Table S5). The Gentry dataset also has a disproportionate number of communities with the opposite tendencies to the other datasets for the other shape metrics– i.e., an overrepresentation of communities with high Simpson’s evenness and Shannon diversity, and low skewness.

**Table 1.** Proportions of extreme values for percentile scores for observed SADs compared to samples from the feasible set. For dissimilarity, this is the proportion of percentile scores >95; by chance, ∼5% of scores should be in this extreme. For all other metrics, this is the proportion <2.5 or >97.5; by chance ∼2.5% of scores should be in either extreme. n refers to the number of communities included for each dataset for each metric. The proportions shown are for the directions of effects observed for most datasets; for the opposite-direction effects, see Table S5.

| Dataset | High dissimilarity | High proportion of rare species | High skew | Low Simpson | Low Shannon |
| --- | --- | --- | --- | --- | --- |
| Breeding Bird Survey | 23%; n = 2773 | 4.5%; n = 2773 | 9%; n = 2773 | 21%; n = 2773 | 23%; n = 2773 |
| FIA | 7.2%; n = 18447 | 1.4%; n = 17410 | 2.8%; n = 17410 | 5.8%; n = 17410 | 5.5%; n = 17410 |
| Gentry | 34%; n = 224 | 0.9%; n = 223 | 11%; n = 223 | 9.9%; n = 223 | 7.6%; n = 223 |
| Mammal Communities | 32%; n = 552 | 13%; n = 511 | 12%; n = 505 | 28%; n = 511 | 30%; n = 511 |
| Misc. Abundance | 59%; n = 494 | 27%; n = 486 | 27%; n = 484 | 53%; n = 486 | 56%; n = 486 |

**Figure 3.**
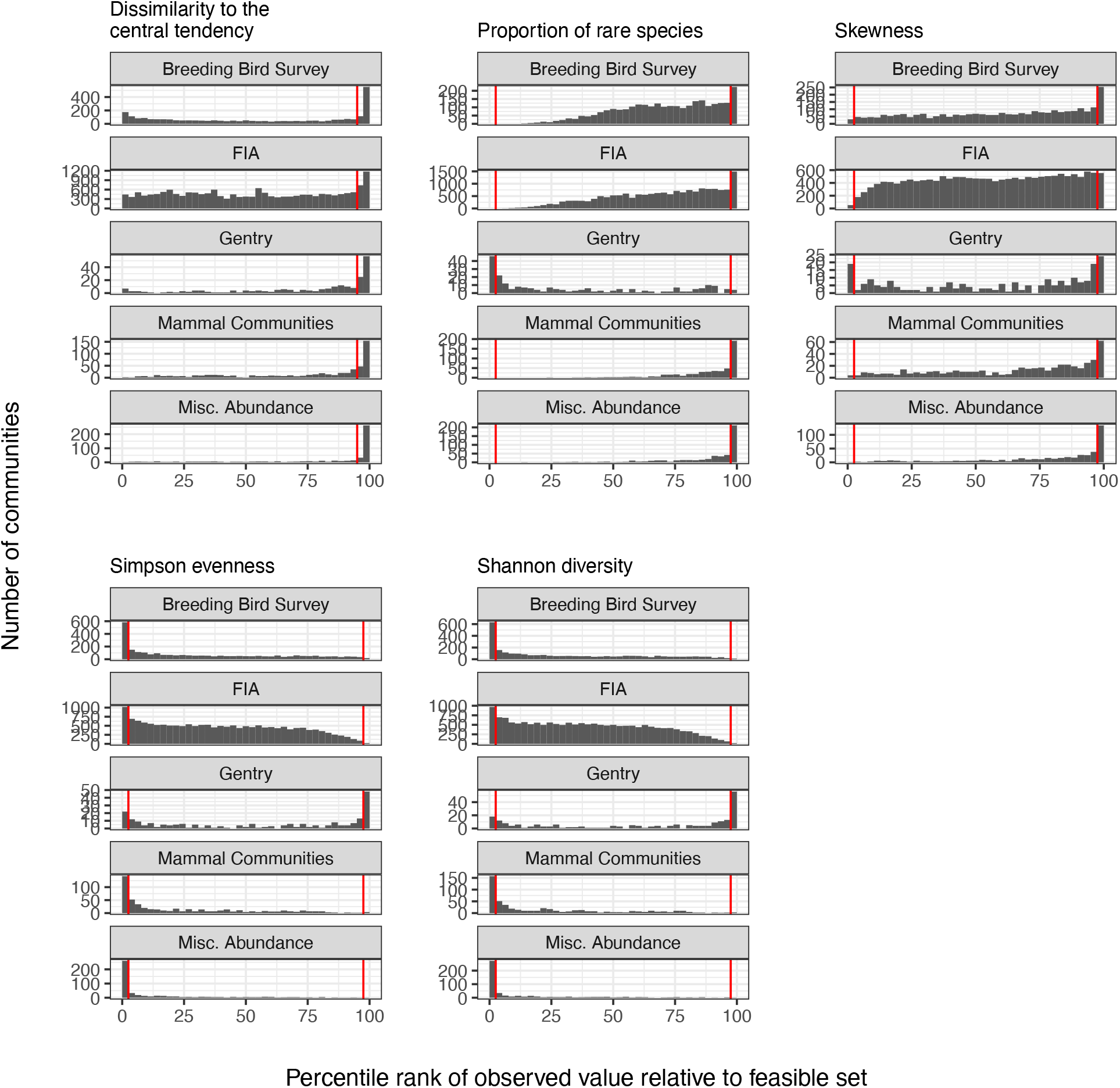
Many ecological communities are highly unusual compared to their statistical baselines. Percentile ranks are calculated by comparing each community to its sampled feasible set, with very high or very low percentile ranks reflecting extreme values relative to statistical baselines. The vertical red lines mark the 95^th^ percentile for the dissimilarity to the central tendency, and the 2.5^th^ and 97.5^th^ percentiles for all other metrics. Species abundance distributions that are sampled at random from the feasible set will produce percentile ranks that are roughly uniformly distributed from 0 to 100, with approximately 5% of values above the 95^th^ percentile or 2.5% of values above and below the 2.5^th^ and 97.5^th^ percentiles, respectively. In contrast, most datasets have more communities that are highly skewed or uneven than would be expected by chance. The percentile values shown are the mean of the percentile scores defined as the proportion of comparison values <=, and <, the focal value. In calculating the proportion of sites with extreme values, the <= designation gives an appropriately conservative estimate of the proportion of high values, but overestimates the proportion of very low values, and the reverse occurs for the < designation.

In contrast to the other datasets, percentile scores for sites from the FIA dataset are more uniformly distributed, and the proportions of extreme values are closer to what would be expected by chance (Figure 3, Table 1). Only 7% of FIA communities are highly dissimilar to their feasible sets (compared to a random expectation of 5%). Among the shape metrics, only 2.7% (compared to 2.5% at random) of sites have high values for skewness, 1.3% have high proportions of rare species, 5.7% have low Simpson’s evenness, and 5.4% have low Shannon diversity.

### The narrowness of the expectation

The ability to detect deviations from the statistical baseline depends in part on the distribution of SADs in the feasible set. Overall, as the size of the feasible set increases, the SADs in a feasible set become more narrowly clustered around the central tendency of that feasible set, and the sampled distributions for shape metrics generally become less variable (Figure 4). In small communities, the breadth indices are highly variable and often very large – approaching 1, meaning that a 95% density interval of the values in the distribution spans nearly the entire range of values – while the breadth indices for larger communities rarely exceed ∼.7 for skewness, Simpson evenness, and Shannon diversity, and ∼.8 for the proportion of rare species. Among our datasets, the FIA and Mammal Community databases have the smallest communities, in terms of S and N, and tend to have the largest proportions of feasible sets with high breadth indices (Figure S6).

**Figure 4.**
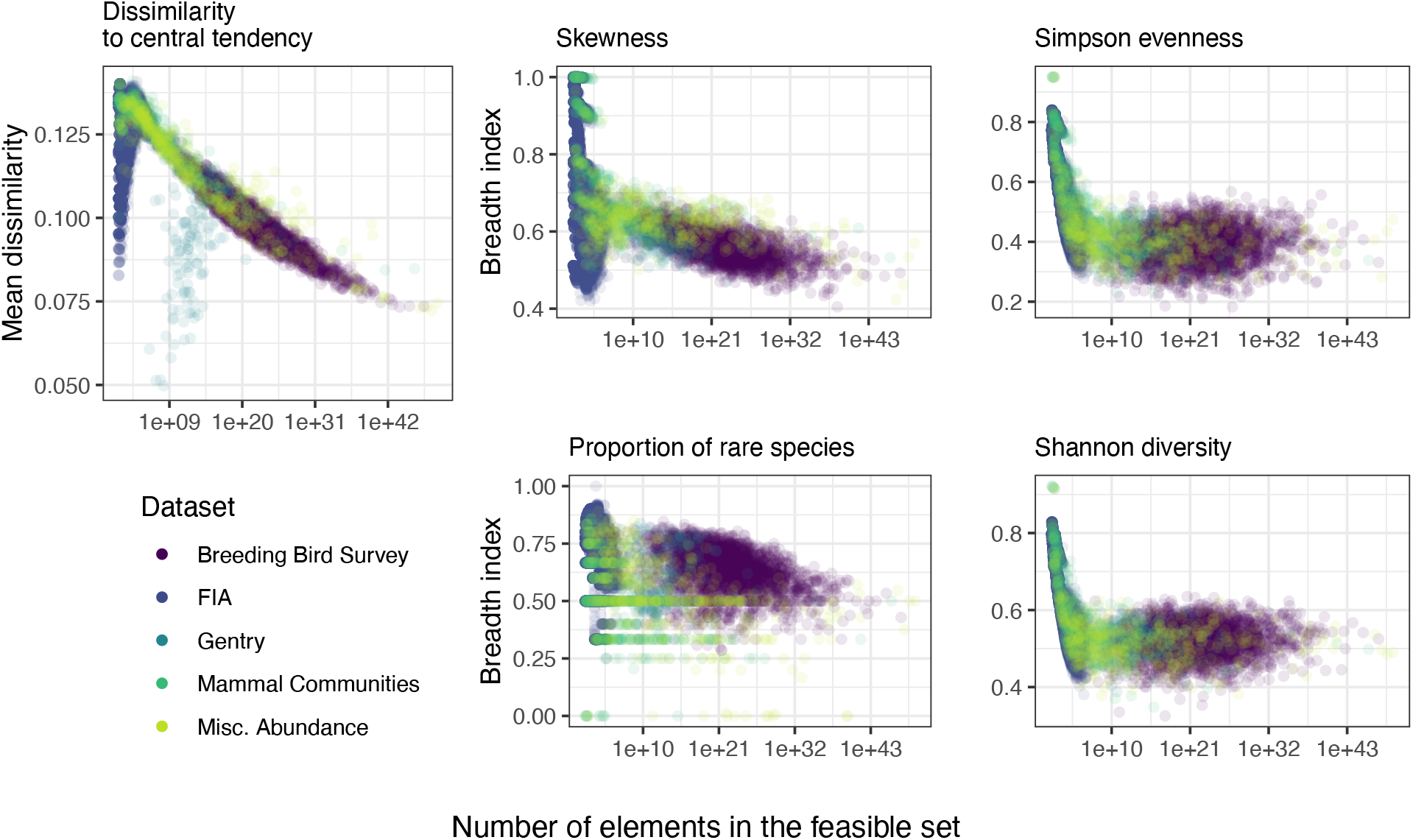
The variability of the feasible set, defined as either the mean dissimilarity of elements of the feasible set to the central tendency of the feasible set, or via a breadth index (see Figure 1), decreases as the number of possible SADs in the feasible set becomes very large. Highly variable feasible sets constitute broad, poorly-defined statistical baselines that may impede our ability to confidently detect deviations between observations and what is expected given the baseline. Small feasible sets, which occur for small combinations of S and N, are often highly variable. The majority of these small, highly variable feasible sets occur for communities in the FIA and Mammal Community databases. Although the Gentry dataset also contains communities with small feasible sets, these communities also have a very low ratio of N to S, meaning their entire feasible sets may be constrained to be more self-similar than small feasible sets in general (see Dissimilarity to central te∂åndency). There is, however, substantial additional variation in the dissimilarity and breadth indices not accounted for by the size of the feasible set or the ratio of N to S.

### Sensitivity to sampling variability

In almost all cases, SADs adjusted for the under-observation of rare species are even more extreme relative to their feasible sets than unadjusted SADs (Figure 5; see Appendix A7 for complete results of resampling). For all datasets, adjusted SADs show more high values for skewness and the proportion of rare species, and low values for Simpson’s evenness and Shannon diversity, than unadjusted SADs.

**Figure 5.**
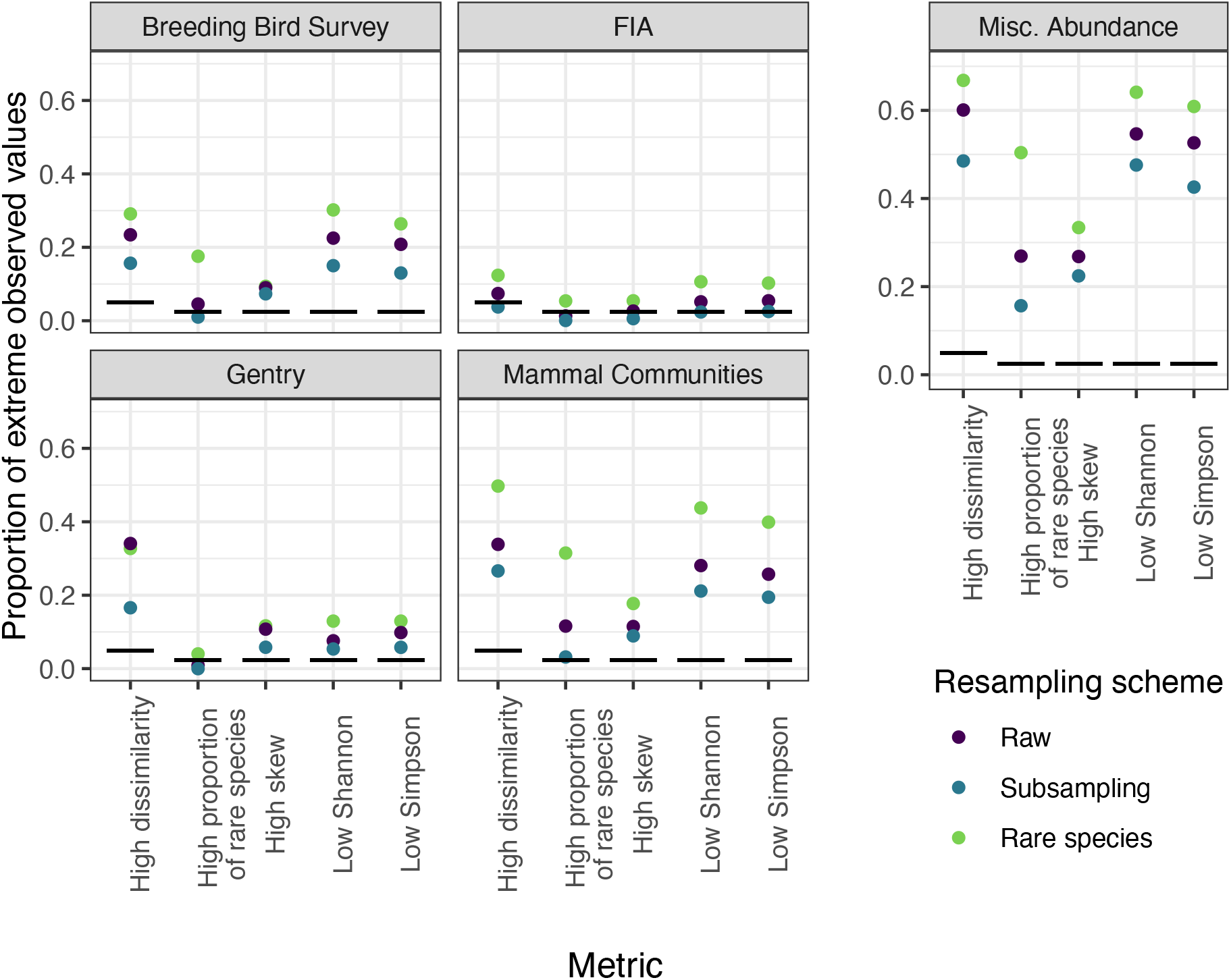
Summaries of how resampling to adjust for under-detection of rare species (green) and subsampling (blue) change the proportion of extreme values observed for each metric and dataset. The horizontal black lines mark the approximate proportions of extreme values that would be expected at random: 5% for dissimilarity to the central tendency, and 2.5% for all other metrics. Adjusting for rare species consistently increases the proportion of extreme values relative to the raw SADs, while subsampling often decreases it but generally does not eliminate or change the direction of the effect. The exception is for the FIA dataset, which does not show strong deviations for either raw or resampled SADs. Shown are the effects and directions observed for most datasets; for complete results of resampling, including the opposite direction effects, see A7.

Subsampling consistently reduces the proportion of extreme observations across all datasets and metrics (Figure 5; Appendix A7). In most instances, the proportion of extreme observations still exceeds the proportion that would be expected by chance. However, the proportion of sites with high numbers of rare species observed for the BBS and Mammal Community databases drop from 4.5% to 1% and ∼13% to 3.5% with resampling. For FIA, the proportions of sites with high dissimilarity, low evenness and Shannon diversity all drop from 6-8% to 2-3%. Note that, for FIA, neither the raw nor the resampled SADs have a disproportionate representation of extreme values for the remaining metrics.

## Discussion

We found widespread evidence that SADs for a range of real ecological communities deviate from the forms expected given the distribution of shapes within their feasible sets. Overall, these deviations may signal that ecological processes operate on top of statistical constraints, thereby driving the SAD away from shapes generated by purely statistical processes. We also found that the magnitude and form of deviation varied among the datasets we considered. This variability may reflect statistical phenomena related to the size of S and N and their ratio, or it may reflect different biological processes dominating in different contexts. Finally, although a disproportionate number of communities deviated statistically from their feasible sets, there were also many communities for which we did not detect deviations. This does not imply the absence of ecological processes operating on these SADs. Rather, one possible explanation is that multiple ecological processes are operating simultaneously and with countervailing effects, resulting in no dominating net impact on the shape of the distribution beyond that imposed by fundamental constraints (Harte 2011; Harte and Newman 2014). Going forward, testing whether ecological theories or common functional approximations (e.g. the log-normal distribution) accurately predict the deviations between observed SADs and their statistical baselines may be much more fruitful than focusing only on the general form of the SAD (McGill et al. 2007; Locey and White 2013; Harte and Newman 2014).

In most cases, and most pronouncedly for the Breeding Bird Survey, Mammal Community, and Miscellaneous Abundance databases, our results suggest that the prevailing processes cause abundance distributions to be highly uneven, rather than those that produce more even abundances across species. For these communities, observed SADs tended to be unusually skewed and uneven, and to have a high proportion of rare species, compared to their feasible sets. Accounting for undersampling of rare species strengthened these effects, while subsampling weakened them. Perhaps unsurprisingly, the effect of these two resampling approaches was especially noticeable for the proportion of rare species; enriching the SAD directly adds rare species, while subsampling is likely to drop rare species even if it otherwise recaptures the general shape of a distribution. The long tail of rare species in the SAD has been a consistent focus in SAD research, and our results highlight that the rare tails of observed SADs are extraordinary, even among the hollow-curve shapes that dominate the feasible set. Ecological processes may lengthen the rare tail and decrease the evenness of the SAD, for example by promoting the persistence of rare species at very low abundances (Yenni et al. 2012). Or, they could drive abundant species to have larger populations than would be statistically expected, without also driving other species entirely to extinction (Chesson 2000).

While the Gentry database also exhibits deviations tending towards high unevenness, an even greater proportion of its communities are *more* even, and have a lower proportion of rare species, than would be expected given their feasible sets. This could indicate that there are biological differences between the systems in the Gentry and other datasets that result in different forms for the SAD. Alternatively, the statistical characteristics of the feasible sets for these communities could modulate the detected deviations. Communities in the Gentry database have high species richness and low average abundance (Figure 1). Among these, many of the communities exhibiting high evenness and low proportions of rare species are those with very high species richness and low average abundance (N/S < ∼3) (see Appendix A8). As a result, these communities have unusual statistical baselines: the corresponding feasible sets have the highest proportions of rare species of any of the feasible sets in our analysis. Although observed SADs for these communities also have high proportions of rare species, taking the statistical baseline into account would suggest that the extraordinary thing about these SADs is that they do not have even more rare species. Simultaneously, there may be biological reasons why the species-rich but relatively low-abundance tropical tree communities of the Gentry database differ from those in other datasets. The same mechanisms that promote high diversity may manifest in high evenness, and/or ecological features particular to these forests may produce unusual shapes for the SAD. Because no communities from our other datasets are comparable in S and N, we cannot disentangle these statistical and biological explanations. This is an excellent opportunity to develop additional theoretical and empirical approaches to predict and explain variation in the deviations between SADs and their feasible sets, in particular for species-rich communities across ecosystems.

Unlike the other four datasets, communities in the FIA dataset showed weak or no evidence of deviations from their feasible sets. We entertained two general classes of explanation for why the FIA dataset differs from the others in our analysis: first, that biological attributes of the FIA communities cause the SADs for these communities to differ from the others in our database, and second, that statistical phenomena related to S and N may modulate the capacity to detect deviations for these communities. To distinguish between possible biological drivers causing FIA to differ from the other datasets, and factors intrinsic to S and N, we compared a subset of ∼300 FIA communities to communities from other datasets with directly matching S and N. We did not find differences in the distribution of percentile scores for any metrics between communities from FIA and communities from other datasets, confirmed via Kolmogorov-Smirnov tests (Appendix A9). Although 300 communities constitute a small sample relative to the 20,355 FIA communities we analyzed, these results point to statistical phenomena, and not biological attributes unique to FIA, as the likely explanation for the differences.

A second possibility is that these differences reflect statistical phenomena related to community size in terms of S, N, and as a result, the number of possible SADs in a community’s feasible set. The FIA communities are the smallest across our datasets (Figure 1), and communities with small values of S and N have smaller feasible sets. When there are relatively few possible SADs in the feasible set, they may be less tightly clustered around their central tendencies, and the distributions for their shape metrics may be less narrowly peaked, than when there are very large numbers of possible SADs. High variability within the feasible set weakens the statistical distinction between “common” and “extreme” shapes (Figure 2). Under these circumstances, any deviations – or lack thereof – will be less informative than for communities with more strongly defined statistical baselines (Jaynes 1957). The average dissimilarity to the central tendency, and the distributions of breath indices for specific metrics, broadly align with this principle. Across the range of community sizes represented in our datasets, small feasible sets have highly variable, and often very broad, feasible sets (Figure 4). More specifically, very small communities – for example, those with fewer than 2000 possible SADs in their feasible sets, or S ∼ 20 and N ∼ 40 – exhibit more highly variable feasible sets than large communities, and these small communities also show less consistent deviations (Figure 6; Appendix A10). Of our datasets, FIA is most dominated by small communities (68% of communities have fewer than 2000 possible SADs), and these small-community phenomena may therefore have the greatest impact on results aggregated over the FIA dataset.

**Figure 6.**
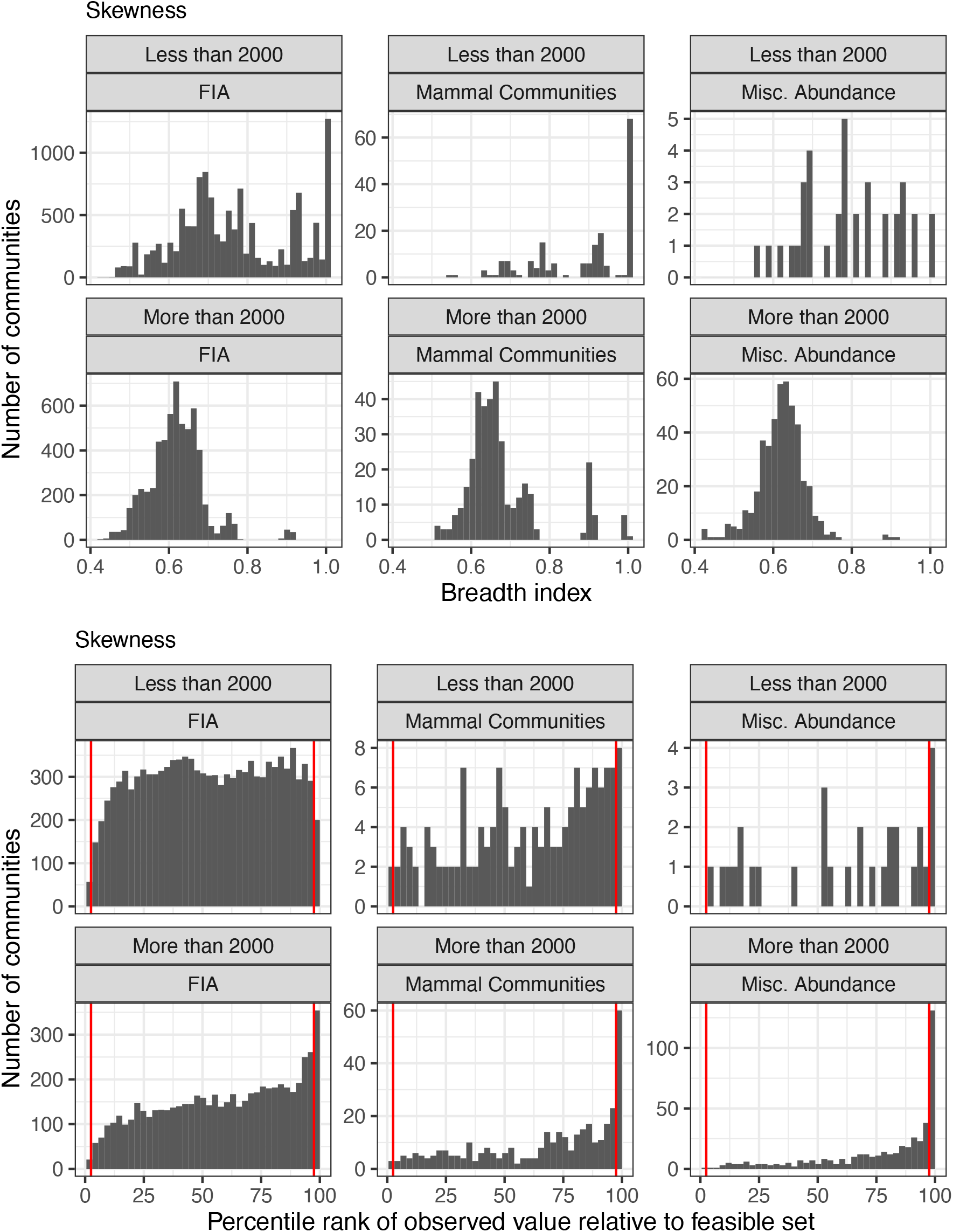
Very small communities (e.g. those with fewer than 2000 possible SADs in the feasible set; upper rows) exhibit more variable, broadly-defined statistical baselines (top) and less consistently extreme observed values relative to their feasible sets (bottom). 2000 possible SADs is used as a cutoff because it allows for a comparison using a substantial number of communities from the FIA and two other datasets. Of these datasets, the FIA is the most dominated by very small communities (68% of FIA sites have fewer than 2000 possible SADs, compared to 34% for the Mammal Community and 7% for the Miscellaneous Abundance databases). Results shown are for skewness; for complete results see Appendix A10.

If it is true that the highly variable feasible sets associated with small communities contribute to the weak evidence of deviations observed for the FIA dataset, such considerations affect our capacity to use this approach to distinguish signal from noise for a substantial contingent of ecological communities. Because the combinations of S and N represented in our analyses are irregularly distributed among different datasets (Figure 1), and because there is a great deal of variation in our breadth indices not accounted for by the size of the feasible set (Figure 4), we do not interpret these results as showing a threshold for defining problematically small communities. A more systematic exploration of the S and N state space, combined with more nuanced metrics for characterizing the variability of the feasible set, could clarify the relationship between S and N, the size of the feasible set, and statistical power. However, FIA and other small, highly variable communities have on the order of 10-20 species and 30-60 individuals, suggesting a general range of values below which we have diminished power to detect deviations from the statistical baseline represented by the feasible set. Communities with on the order of 5 species, or 100s to 1000s of individuals, have previously been suggested as “small” in this context (Preston 1948; McGill et al. 2007). To meaningfully draw inferences using deviations in these small communities, we will need more sensitive metrics than those used here, and/or theories that generate more specific predictions for the SAD. In the absence of such, we may stand to learn the most by focusing on SADs from relatively large communities.

It is also important to recognize that there are multiple plausible approaches to defining a statistical baseline for the SAD, of which we have taken only one (Haegeman and Loreau 2008, Locey and White 2013). Our approach follows Locey and White (2013) and reflects the random partitioning of individuals into species, with the resulting distributions considered unique if the species’ abundance values are unique, regardless of the order in which the values occur. This philosophy reflects a longstanding approach in the study of abundance distributions: to focus on the shape of the distribution without regard to species’ identities (McGill et al 2007). Other assumptions regarding the statistical baseline may be equally valid and generate different statistical expectations, which may alter if, and in what ways, empirical distributions appear unusual. For example, incorporating differences in species order into the statistical baseline – which would imply that identifying *which* species contain the most or least individuals is important – might reduce the representation of long-tailed, highly uneven SADs within the feasible set, and make the rare tail observed for real SADs appear more unlikely than it does here. Under our assumptions, the SADs (1,2,3,4) and (1, 1, 1, 7) each count as only one unique SAD. Taking species order into account would mean that (1,2,3,4) would count as 24 (4!) unique SADs, because there are 4! ways to assign the abundances to each species. However, an SAD containing species with equal abundances, such as (1, 1, 1, 7), would only count as 4 unique SADs. For SADs, equal abundances are likely most prevalent among rare species. If this is true, then this set of assumptions would generate feasible sets where rare-tailed SADs are relatively scarce, making observed SADs with rare tails seem even more extraordinary. Additional formulations for the statistical baseline exist, including those that approximate exponential, Poisson, or log-series distributions in the limit (Harte et al. 2008, Favretti 2018). Investigating and comparing the results that emerge from different baselines will be an important next step towards reinvigorating the use of the SAD as a diagnostic tool.

Our study demonstrates the utility, and the potential challenges, of applying tools from the study of complex systems and statistical mechanics to the study of ecological communities (Haegeman and Loreau 2008, Harte 2011, White et al. 2012, Harte and Newman 2014). While concepts such as maximum entropy and the feasible set are promising horizons for macroecology, the small size of some ecological communities may present difficulties that are rare in the domains for which these tools were originally developed (Jaynes 1957, Haegeman and Loreau 2008). When the observed numbers of species and individuals are too small to generate highly resolved statistical baselines, these approaches will be less informative than we might hope – as appears to be the case for the smallest communities in our analysis. In larger communities, where mathematical constraints have more resolved effects on the form of the SAD, our results show that these constraints alone do not fully account for the extremely uneven SADs we often observe in nature – leaving an important role for ecological processes. This ability to detect and diagnose the specific ways in which empirical SADs deviate from randomness can generate new avenues for understanding how and when biological drivers affect the SAD. There are, of course, still many elements to be improved in our ability to distinguish biological signal from randomness, including assessing alternative statistical baselines and calibrating our expected power to detect deviations, especially for small communities. Indeed, more sensitive metrics could also enable identification of processes that operate through time. Continuing to explore and account for the interplay between statistical constraint and biological process constitutes a promising and profound new approach to our understanding of this familiar, yet surprisingly mysterious, ecological pattern.

## Acknowledgements

RMD was supported by the National Science Foundation Graduate Research Fellowship under Grant No. DGE-1315138 and DGE-1842473. HY’s time was supported by Gordon and Betty Moore Foundation’s Data-Driven Discovery Initiative, Grant GBMF4563, awarded to Ethan White. We thank Erica Newman, Justin Kitzes, and Ethan White for helpful and illuminating discussions.

