## Appendix S1 for "Empirical abundance distributions are more uneven than expected given their statistical baseline"

Appendix S1 – Details of filtering datasets

We filter datasets in two stages. First, we filter **prior to trying to sample from the feasible set**, in order to remove communities that have a combined S and N too high for us to sample the feasible set, and to reduce the Forest Inventory and Analysis dataset to a manageable size. Second, we filter **after sampling the feasible but before we aggregate results across communities**, to remove cases where mathematical constraints result in uninformative results. Below we elaborate on the logic and methods for these filtering processes, including code to allow others to see precisely what was implemented.

### Pre-feasible set sampling

While our algorithm for sampling the feasible set improved our ability to assess the shape of the feasible set for larger communities, there were still computational limits on what we could do. We filtered out all communities with more than 40720 individuals, because this is the largest community we were able to sample given the available resources. This upper limit results in the removal of a total of 4 communities, all of them from the Miscellaneous Abundance Database.

For computational reasons, we also created a sub-sample of the FIA dataset, because sampling the feasible set for all 103,343 FIA communities overwhelms our computational pipeline. Because the FIA has so many extremely small communities (92,988 with fewer than 10 species), we decided to randomly select a sample of the small communities for analysis. Thus our FIA dataset consisted of ~20 ,000 communities comprising all communities with more than 10 species (10,355) plus 10,000 randomly selected communities with 3-10 species.

Code for downloading the data (and all other analyses) used for this project can be found at www.github.com/diazrenata/scadsanalysis. The data download functions are at <https://github.com/diazrenata/scadsanalysis/blob/master/R/download_data.R>.

#### Overview

The download_data function downloads raw data files from <https://github.com/weecology/sad-comparison/> (for BBS, Gentry, Mammal Community Database, and FIA; data from Baldridge (2016) and also used in White, Thibault, and Xiao (2012)) and figshare <http://figshare.com/files/3097079> (for the Miscellaneous Abundance Database; Baldridge (2015)). These raw files are stored in working-data\abund_data and are not edited.

The datasets downloaded from GitHub exactly match the corresponding data files in the Zenodo archive for Baldridge (2016), available here: (https://zenodo.org/record/166725). If in the future it is preferred or necessary to use the archival versions, they can be downloaded and stored in working-data\abund_data and the rest of the analysis should run just as if they had been downloaded from GitHub.

The Miscellaneous Abundance and FIA databases undergo additional filtering steps, implemented in the filter_misc_abund and filter_fia_short and filter_fia_small functions. These functions run automatically when the data are downloaded and save new files to working-data\abund_data.

The load_dataset function loads either the filtered datasets (if specified, for Misc. Abund and FIA) or the raw data (for all others, or if specified for Misc. Abund and FIA), and adds additional columns and column types to prepare the data for the computational pipeline. The datasets as returned from load_dataset are exactly what goes into the analysis.

In the following sections, we load the raw data files (as they were downloaded), perform the filtering process manually (if applicable), and compare the resulting data to the data that are used in the analysis, i.e. the data that we get if we use load_dataset.

#### BBS (no filtering)

For BBS, and all datasets that are not filtered, we can simply load the data from the file downloaded by download_data, add the columns and column formats that are added by load_data, and compare the result to the result of load_data:

bbs_raw <- read.csv(here::here("working-data", "abund_data", "bbs_spab.csv"), stringsAsFactors = F, header = F, skip = 2)

colnames(bbs_raw) <- c("site", "year", "species", "abund")

bbs_raw <- bbs_raw %>%
 mutate(site = as.character(site),
 dat = "bbs",
 singletons = F,
 sim = -99,
 source = "observed") %>%
 group_by(site) %>%
 arrange(abund) %>%
 mutate(rank = row_number()) %>%
 ungroup()

bbs_loaded <- load_dataset("bbs")

Here we confirm that all values for abund and site match between the version we get from load_dataset and the version we just created from the raw data file:

any(!(bbs_loaded$abund == bbs_raw$abund))

\## [1] FALSE

any(!(bbs_loaded$site == bbs_raw$site))

\## [1] FALSE

Here we check to confirm that no communities have a total abundance exceeding our upper limit (40720, marked by the horizontal red line):

bbs_statevars <- get_statevars(bbs_raw)

ggplot(bbs_statevars, aes(s0, n0)) +
 geom_point() +
 theme_bw() +
 geom_hline(yintercept = 40720, color = "red")

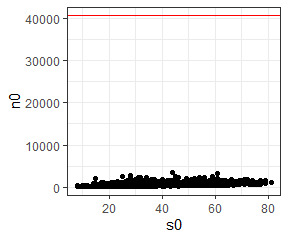

No communities in BBS have more than 40720 individuals, so all are initially included in the analysis.

#### Gentry (no filtering)

For Gentry, as with BBS, we can simply load the data from the file downloaded by download_data, add the columns and column formats that are added by load_data, and compare the result to the result of load_data:

gentry_raw <- read.csv(here::here("working-data", "abund_data", "gentry_spab.csv"), stringsAsFactors = F, header = F, skip = 2)

colnames(gentry_raw) <- c("site", "year", "species", "abund")

gentry_raw <- gentry_raw %>%
 mutate(site = as.character(site),
 dat = "gentry",
 singletons = F,
 sim = -99,
 source = "observed") %>%
 group_by(site) %>%
 arrange(abund) %>%
 mutate(rank = row_number()) %>%
 ungroup()

gentry_loaded <- load_dataset("gentry")

Here we confirm that all values for abund and site match between the version we get from load_dataset and the version we just created from the raw data file:

any(!(gentry_loaded$abund == gentry_raw$abund))

\## [1] FALSE

any(!(gentry_loaded$site == gentry_raw$site))

\## [1] FALSE

Here we check to confirm that no communities have a total abundance exceeding our upper limit (40720, marked by the horizontal red line):

gentry_statevars <- get_statevars(gentry_raw)

ggplot(gentry_statevars, aes(s0, n0)) +
 geom_point() +
 theme_bw() +
 geom_hline(yintercept = 40720, color = "red")

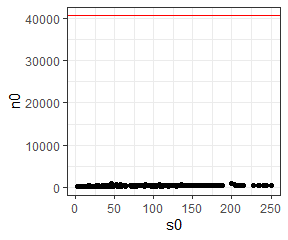

No communities in Gentry have more than 40720 individuals, so all are initially included in the analysis.

#### Mammal Community Database (not filtered)

For MCDB, as with BBS and Gentry, we can simply load the data from the file downloaded by download_data, add the columns and column formats that are added by load_data, and compare the result to the result of load_data:

mcdb_raw <- read.csv(here::here("working-data", "abund_data", "mcdb_spab.csv"), stringsAsFactors = F, header = F, skip = 2)

colnames(mcdb_raw) <- c("site", "year", "species", "abund")

mcdb_raw <- mcdb_raw %>%
 mutate(site = as.character(site),
 dat = "mcdb",
 singletons = F,
 sim = -99,
 source = "observed") %>%
 group_by(site) %>%
 arrange(abund) %>%
 mutate(rank = row_number()) %>%
 ungroup()

mcdb_loaded <- load_dataset("mcdb")

Here we confirm that all values for abund and site match between the version we get from load_dataset and the version we just created from the raw data file:

any(!(mcdb_loaded$abund == mcdb_raw$abund))

\## [1] FALSE

any(!(mcdb_loaded$site == mcdb_raw$site))

\## [1] FALSE

Here we check to confirm that no communities have a total abundance exceeding our upper limit (40720, marked by the horizontal red line):

mcdb_statevars <- get_statevars(mcdb_raw)

ggplot(mcdb_statevars, aes(s0, n0)) +
 geom_point() +
 theme_bw() +
 geom_hline(yintercept = 40720, color = "red")

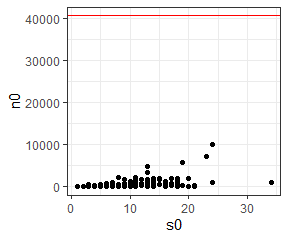

No communities in the MCDB have more than 40720 individuals, so all are initially included in the analysis.

#### Miscellaneous Abundance Database (filtered)

For the Miscellaneous Abundance Database, we can proceed similarly to with the non-filtered datasets. We load the data from the file downloaded by download_data, add the columns and column formats that are added by load_data, and compare the result to the result of load_data.

However, Misc. Abund includes datasets reported as relative abundance in addition to count data. Relative abundance data is not appropriate for this analysis, so we filter out records where abund = 0 to restrict our analysis to communities with count data. Also, because we will eventually filter out some highly abundant communities, at this stage we make sure to load the **unfiltered** version using load_dataset.

misc_abund_raw <- read.csv(here::here("working-data", "abund_data", "misc_abund_spab.csv"))

misc_abund_raw <- misc_abund_raw %>%
 dplyr::rename(site = Site_ID,
 abund = Abundance)

misc_abund_raw <- misc_abund_raw %>%
 mutate(site = as.character(site),
 dat = "misc_abund",
 singletons = F,
 sim = -99,
 source = "observed") %>%
 filter(abund > 0) %>%
 group_by(site) %>%
 arrange(abund) %>%
 mutate(rank = row_number()) %>%
 ungroup()

misc_abund_loaded <- load_dataset("misc_abund")

Here we confirm that, if we load the unfiltered dataset using load_dataset, the records match the version we have created manually from the raw file:

any(!(misc_abund_loaded$abund == misc_abund_raw$abund))

\## [1] FALSE

any(!(misc_abund_loaded$site == misc_abund_raw$site))

\## [1] FALSE

Here we check whether any communities have more than 40720 individuals (horizontal red line):

misc_abund_statevars <- get_statevars(misc_abund_raw)

ggplot(misc_abund_statevars, aes(s0, n0)) +
 geom_point() +
 theme_bw() +
 geom_hline(yintercept = 40720, color = "red")

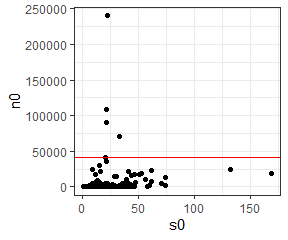

Misc abund has 4 communities with more than 40720 individuals, and these communities are removed:

misc_abund_sv_filtered <- misc_abund_statevars %>%
 filter(n0 <= 40720)

misc_abund_filtered <- filter(misc_abund_raw, site %in% misc_abund_sv_filtered$site)

We can access the **filtered** version of the dataset, as used in the analysis, using load_dataset("misc_abund_short"). We then check to confirm that the filtered version of the dataset that we get from load_dataset matches the version that we filtered manually:

misc_abund_short_loaded <- load_dataset("misc_abund_short")

any(!(misc_abund_short_loaded$abund == misc_abund_filtered$abund))

\## [1] FALSE

any(!(misc_abund_short_loaded$site == misc_abund_filtered$site))

\## [1] FALSE

#### FIA (filtered)

Again, we can begin similarly to with the non-filtered datasets. We load the data from the file downloaded by download_data, add the columns and column formats that are added by load_data, and compare the result to the result of load_data. Because we will remove some communities from FIA, we make sure to load the **unfiltered** version using load_data.

fia_raw <- read.csv(here::here("working-data", "abund_data", "fia_spab.csv"), stringsAsFactors = F, header = F, skip = 2)

colnames(fia_raw) <- c("site", "year", "species", "abund")

fia_raw <- fia_raw %>%
 mutate(site = as.character(site),
 dat = "fia",
 singletons = F,
 sim = -99,
 source = "observed") %>%
 filter(abund > 0) %>%
 group_by(site) %>%
 arrange(abund) %>%
 mutate(rank = row_number()) %>%
 ungroup()

fia_loaded <- load_dataset("fia")

Here we confirm that, if we load the unfiltered dataset using load_dataset, the records match the version we have created manually from the raw file:

any(!(fia_loaded$abund == fia_raw$abund))

\## [1] FALSE

any(!(fia_loaded$site == fia_raw$site))

\## [1] FALSE

Here we check whether any communities have more than 40720 individuals (horizontal red line):

fia_statevars <- get_statevars(fia_raw)
ggplot(fia_statevars, aes(s0, n0)) +
 geom_point() +
 theme_bw() +
 geom_hline(yintercept = 40720, color = "red")

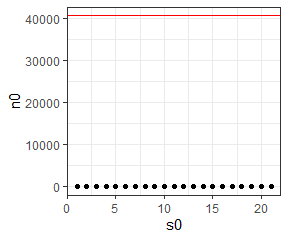

FIA has no communities with more than 40720 individuals. However, it presents additional issues. It has 103343 communities, of which 92988 have fewer than 10 species (vertical red line in the figure below)

ggplot(fia_statevars, aes(s0, n0)) +
 geom_point() +
 theme_bw() +
 geom_vline(xintercept = c(9.5), color = "red")

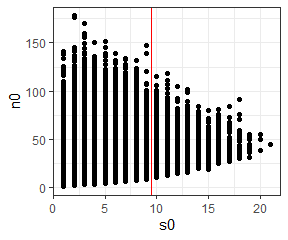

This many communities overwhelms our computational pipeline. We therefore sample all 10355 communities with 10 or more species, and a random subsample of 10,000 communities with 3-9 species. We then run these through the pipeline as two separate databases. fia_short is the communities with 10 or more species, and fia_small is the 10,000 communities with 3-9 species. We re-combine them as “FIA” for aggregate analyses. This results in a total of 20,355 FIA communities in the analysis.

fia_sv_short <- fia_statevars %>%
 dplyr::filter(s0 >= 10)

fia_short <- fia_raw %>%
 dplyr::filter(site %in% fia_sv_short$site) %>%
 dplyr::mutate(dat = "fia_short")

fia_short_statevars <- get_statevars(fia_short)

ggplot(fia_short_statevars, aes(s0, n0)) +
 geom_point() +
 ggtitle("fia short, >= 10 species") +
 theme_bw()

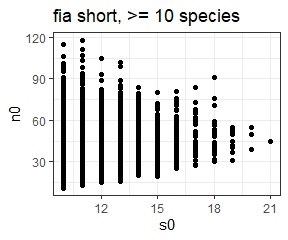

fia_sv_small <- fia_statevars %>%
 dplyr::filter(s0 >= 3) %>%
 dplyr::filter(s0 <= 9)

 set.seed(1977)
 fia_sv_small <- fia_sv_small[ sample.int(nrow(fia_sv_small), size = 10000, replace = F), ]


fia_small <- fia_raw %>%
 dplyr::filter(site %in% fia_sv_small$site) %>%
 dplyr::mutate(dat = "fia_small")

fia_small_statevars <- get_statevars(fia_small)

ggplot(fia_small_statevars, aes(s0, n0)) +
 geom_point() +
 ggtitle("fia small, 3-9 species")

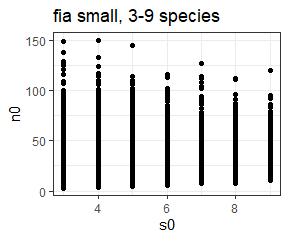

The “short” and “small” datasets are saved as .csvs and can be loaded using load_dataset as follows. These are the versions that are used in the analysis.

fia_small_loaded <- load_dataset("fia_small")

fia_short_loaded <- load_dataset("fia_short")

Here we confirm that versions of “fia_short” and “fia_small” that we created manually match the ones we obtain from load_dataset:

any(!(fia_small_loaded$abund == fia_small$abund))

\## [1] FALSE

any(!(fia_small_loaded$site == fia_small$site))

\## [1] FALSE

any(!(fia_short_loaded$abund == fia_short$abund))

\## [1] FALSE

any(!(fia_short_loaded$site == fia_short$site))

\## [1] FALSE

### Post-sampling

Some communities fall into mathematically special spaces where basic constraints prevent informative results. We removed all communities that fell into any of the below cases:

1) When communities have only 1 mathematically possible SAD (S = 1, N = S, or N = S + 1).
2) When the sampled feasible sets yield fewer than 20 unique values for skewness/evenness. If there are fewer than 20 unique values in the comparison vector, it is impossible to be in the 5th or 95th percentile.
3) When a community consists of only 2 species, we excluded it from analyses using skewness, because e1071::skewness() always evaluates to 0 if S = 2.

Below is the procedure for performing this filtering (after sampling the feasible set). First, we load our results from all_di.csv. This file contains the results of comparing the skewness and evenness for **observed** SADs to the distributions of skewness and evenness obtained from their sampled feasible sets; “all_di” stands for “all diversity indices”. Every community that was included for sampling is included in all_di. (Additionally, every community has a “singletons” counterpart, which is the same analysis run adjusted for rarefaction. That analysis is discussed elsewhere, and we ignore the rarefaction-adjusted versions here).

all_statevars <- bind_rows(bbs_statevars, fia_short_statevars, fia_small_statevars, gentry_statevars, mcdb_statevars, misc_abund_sv_filtered)


all_di <- read.csv(here::here("analysis", "reports", "all_di.csv"), stringsAsFactors = F)

all_di <- all_di %>%
 filter(!singletons) %>%
 mutate(dat = ifelse(grepl(dat, pattern = "fia"), "fia", dat),
 dat = ifelse(dat == "misc_abund_short", "misc_abund", dat)) %>%
 mutate(Dataset = dat,
 Dataset = ifelse(Dataset == "fia", "Forest Inventory and Analysis", Dataset),
 Dataset = ifelse(Dataset == "bbs", "Breeding Bird Survey", Dataset),
 Dataset = ifelse(Dataset == "mcdb", "Mammal Community DB", Dataset),
 Dataset = ifelse(Dataset == "gentry", "Gentry", Dataset),
 Dataset = ifelse(Dataset == "misc_abund", "Miscellaneous Abundance DB", Dataset))

#head(all_di)

Here we confirm that we have the same number of communities in all_di as we have in our set of communities included for sampling.

nrow(all_di) == nrow(all_statevars)

\## [1] TRUE

Here we remove communities for the first case, only one possible SAD (N = S, N = S + 1, or S = 1).

all_di %>%
 group_by_all() %>%
 mutate(only_one_sad = sum(s0 == n0, s0 == 1, n0 == (s0 + 1)) > 0) %>%
 ungroup() %>%
 group_by(dat) %>%
 summarize(total_only_one_sad = sum(only_one_sad),
 total_sites = dplyr::n()) %>%
 ungroup() %>%
 mutate(all_sites_one_sad = sum(total_only_one_sad),
 all_sites = sum(total_sites))

| dat | total_only_one_sad | total_sites | all_sites_one_sad | all_sites |
| --- | --- | --- | --- | --- |
| bbs | 0 | 2773 | 258 | 24647 |
| fia | 176 | 20355 | 258 | 24647 |
| gentry | 0 | 224 | 258 | 24647 |
| mcdb | 56 | 730 | 258 | 24647 |
| misc_abund | 26 | 565 | 258 | 24647 |

all_di_filtered <- all_di %>%
 filter(s0 != n0,
 s0 != 1,
 n0 != (s0 + 1))

nrow(all_di_filtered)

\## [1] 24389

Removing those with only one SAD results in the removal of 258 sites total. 176 from FIA, 56 from MCDB, and 26 from Misc. Abund.

Finally, we filter for cases 2 and 3. That is, for aggregate analyses, we will restrict our analyses to sites whose feasible sets have more than 20 unique values for skewness or evenness (whichever metric we are focused on). For analyses with skewness, we will also remove sites with only 2 species. This results in these final total numbers of communities included for skewness and evenness.

For skewness:

all_di_filtered %>%
 filter(skew_unique > 20, s0 > 2) %>%
 group_by(dat) %>%
 summarize(sites_for_skewness = dplyr::n()) %>%
 mutate(total_sites_for_skewness = sum(sites_for_skewness))

| dat | sites_for_skewness | total_sites_for_skewness |
| --- | --- | --- |
| bbs | 2773 | 22325 |
| fia | 18300 | 22325 |
| gentry | 223 | 22325 |
| mcdb | 537 | 22325 |
| misc_abund | 492 | 22325 |

For evenness:

all_di_filtered %>%
 filter(simpson_unique > 20) %>%
 group_by(dat) %>%
 summarize(sites_for_evenness = dplyr::n()) %>%
 ungroup() %>%
 mutate(total_sites_for_evenness = sum(sites_for_evenness))

| dat | sites_for_evenness | total_sites_for_evenness |
| --- | --- | --- |
| bbs | 2773 | 22142 |
| fia | 18113 | 22142 |
| gentry | 224 | 22142 |
| mcdb | 542 | 22142 |
| misc_abund | 490 | 22142 |
