## Appendix S2 for "Empirical abundance distributions are more uneven than expected given their statistical baseline"

Appendix S2 – Sampling the Space of Feasible Sets

### Species Abundance Distribution (SAD) Definition

For $n\geq s\geq1$, we define a SAD as:

$$\text{x}=\left( x_{1},x_{2},\ldots,x_{s} \right)$$

where

$$\begin{matrix} x_{i}\in\mathbb{Z}^{+} \\ x_{i}\leq x_{j}\text{ for }i\leq j \\ \sum x_{i}=n \end{matrix}$$

In effect, a SAD is a division of $n$ individuals among $s$ classes (i.e. taxonomic categories), and we are interested only in the sizes of those classes, disregarding their specific identity (i.e. the classes are interchangeable).

Note that we deviate slightly from ecological convention in that we order the sizes of the classes in ascending, as opposed to descending order.

### Feasible Sets

A **Feasible Set**, $F(s,n)$, is the set of all possible SADs for a particular $s$ and $n$, and $f(s,n)=|F(s,n)|$ is the cardinality of $F(s,n)$ (i.e. the size of the Feasible Set). The values of $f(s,n)$ are identical to the number of partitions of $n$ into exactly $s$ parts^[1]^.

#### Alternative Representation

First, we describe an alternative representation for SADs, where we describe the relative differences in sizes of consecutive classes instead of the sizes of the classes directly:

$$\mathbf{y}=\left( y_{1},y_{2},\ldots,y_{s} \right)$$

where

$$\begin{matrix} y_{1} & =x_{1} \\ y_{i} & =x_{i}-x_{i-1}\geq0\text{ for }i\geq2 \end{matrix}$$

Thus,

$$\begin{matrix} x_{1} & =y_{1} \\ x_{2} & =y_{1}+y_{2} \\ x_{3} & =y_{1}+y_{2}+y_{3} \\ & \vdots\\ x_{i} & =\sum_{j=1}^{j=i} y_{j} \end{matrix}$$

and

$$\begin{matrix} n & =x_{1}+x_{2}+\ldots+x_{s} \\ & =s\cdot y_{1}+(s-1)\cdot y_{2}+\ldots+1\cdot y_{s} \end{matrix}$$

#### Generating SADs

We describe a generative approach for creating a SAD.

1. Set $i=1$, the index of the class whose size will be determined next. Set $n_{r}=n$, the number of individuals remaining to be allocated. Set $s_{r}=s$, the number of classes reamining to be filled.
2. Choose a value for $y_{1}$. The possible values are $[1,\lfloor\frac{n_{r}}{s_{r}}\rfloor]$.
3. Compute $n_{r}$, the remaining un-allocated individuals, as $n_{r}=n_{r}-s_{r}\cdot y_{i}$. Compute $s_{r}$, the remaining un-allocated classes, as $s_{r}=s_{r}-1$. Increment $i=i+1$.
4. Choose a value for $y_{i}$. The possible values are $[0,\lfloor\frac{n_{r}}{s_{r}}\rfloor]$.
5. Repeat steps 3 and 4, until the SAD is fully generated.

#### Counting Feasible Sets

Mirroring the generative approach, we can define $f(s,n)$ recursively.

1. Define $r(s,n)$ to be the number of ways of allocating $n$ individuals among $s$ classes, and allowing the sizes of the classes to include 0.
2. $f(s,n)=\sum_{y_{1}=1}^{y_{1}=\lfloor\frac{n}{s}\rfloor} r(s-1,n-s\cdot y_{1})$
3. $r(s,n)=\sum_{y_{i}=0}^{y_{i}=\lfloor\frac{n}{s}\rfloor} r(s-1,n-s\cdot y_{i})$

where $r(1,n)=1$ (there is only way to allocate $n$ individuals among 1 class).

#### Sampling Feasible Sets

We can also sample $F(s,n)$ uniformly, using a similarly recursive approach.

1. For all $y_{1}\in[1,\lfloor\frac{n}{s}\rfloor]$, compute $r(s-1,n-s\cdot y_{1})$.
2. The sum of these values of $r$ determine the relative probabilities for the corresponding value of $y_{1}$.
3. Choose $y_{1}$ based on the corresponding weights, $\frac{r(s-1,n-s\cdot y_{1})}{\sum_{y_{1}} r(s-1,n-s\cdot y_{1})}$
4. Choose the remaining $y_{i}$ according to the corresponding values of $r(s,n)$ as used to count the feasible set.
