## Appendix S3 for "Empirical abundance distributions are more uneven than expected given their statistical baseline"

Appendix S3 - Self-similarity of the elements of the feasible set

We are interested in how the self-similarity of the elements of the feasible set varies over gradients in S and N. The more self-similar the feasible set is, the less likely it is for us to obtain a SAD that differs from the majority of the feasible set by chance. The intuition from “common-sense” probability theory, and a common theme in statistical mechanics (Jaynes 1957) is that as the number of possible arrangements becomes large, the majority of possible arrangements should become very similar to each other in broad-scale characteristics. However, we don’t know that this is the case for SADs, nor do we know the values for S and N where this increase in self-similarity begins to occur.

In the manuscript, we use a breadth index defined as ratio of the 95% interval of summary statistic values : the full range of summary statistic values. This can be calculated quickly and is easily interpretable. It also directly reflects the distributions to which we are comparing our observations. However, it is not a widely-used metric. Also, if the summary statistic values are idiosyncratic (skewness in particular can behave counterintuitively), it can reflect those idiosyncrasies.

With more computing, we can explore how similar the elements of the feasible set are to each other by comparing them to each other directly. Then we can ask whether, as the feasible set gets large, the elements become more similar and converge on a dominant form.

### Overview

We can describe how self-similar the elements of a feasible set are via numerous pairwise comparisons. Given a body of samples drawn from a feasible set, we draw two samples and compute some metric that describes how similar these two samples are to each other. We do this many times, making many pairwise comparisons, to generate a distribution of the self-similarity metric for that feasible set. We then compare how self-similar different feasible sets are by comparing the distributions of self-similarity metrics for the different feasible sets.

Here we demonstrate this process for an example feasible set, and then present results for feasible sets spanning the range of S and N present in our data.

### Example

Here we have a bank of 3870 samples from the feasible set for SADs with 7 species and 71 individuals. We draw two of these samples to compare.

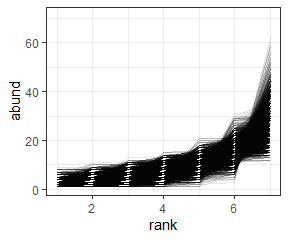

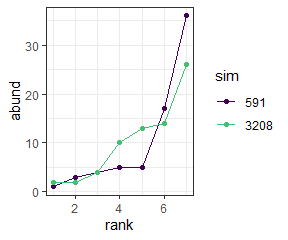

We have implemented 5 metrics of similarity for comparing samples:

- R2 (r2). High R2 indicates higher self-similarity.
- R2 on log-transformed abundances (r2_log). Higher values indicate higher self-similarity.
- The coefficient of determination from a linear model fitting one sample to the other (cd). Higher values indicate higher self-similarity.
- The proportion of individuals allocated to species of differing abundances (prop_off). **Lower** values indicate greater self-similarity.
- The estimated Kullback–Leibler divergence between the two samples (div). **Lower** values indicate greater self-similarity.

| sim1 | sim2 | r2 | r2_log | cd | prop_off | div | s0 | n0 | nparts |
| --- | --- | --- | --- | --- | --- | --- | --- | --- | --- |
| 591 | 3208 | 0.7874279 | 0.7336329 | 0.8397613 | 0.1971831 | 0.1027579 | 7 | 71 | 60289 |

We draw numerous pairs of SADs and compare them numerous times to get distributions for the self similarity metrics.

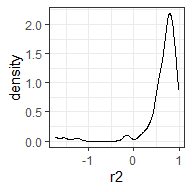

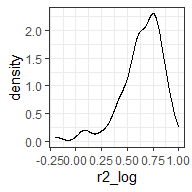

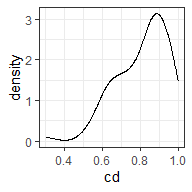

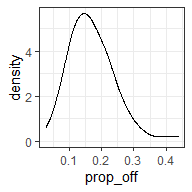

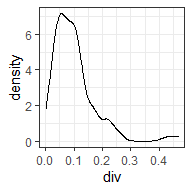

We repeat this process for feasible sets for different values of S and N, and compare the distributions for the self-similarity metrics for the different feasible sets. Here we compare the distributions for the self-similarity metrics for our original feasible set (S = 7, N = 71, “small” in the figure) and for a much larger community (S = 44, N = 13360, “large”).

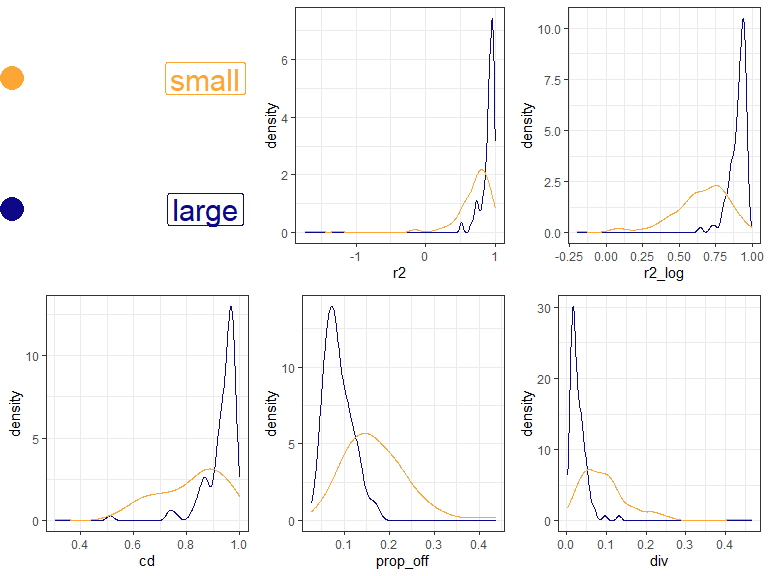

For each metric, the large community is more self-similar than the smaller. That is, it has consistently higher R2, r2_log, and coefficient of determination values, and lower prop_off and K-L divergence.

### Across a range of S and N

Here we have drawn samples from a set of points in S and N space that spans the range present in our datasets. For each feasible set we make 200 comparisons of elements (although for small feasible sets, 200 is not necessarily possible). Here is how that set is distributed in S and N space, colored by the log() number of elements in the feasible set.

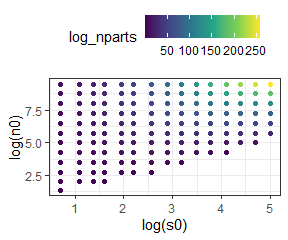

#### Heat maps

We can make heat maps of the density distribution of self-similarity metrics and look at how the densities shift over gradients in the size of the feasible set.

There are a couple of nuances to doing this:

- We need to have the same number of comparisons for every feasible set, so that the density distributions will be comparable. In order to get a large swatch of S by N space, we have taken 50 comparisons from every feasible set. This is pretty low, so it lets us get even small feasible sets. Setting the minimum higher does not change the overall impression.
- The metrics vary in how they are bounded. For some of them (R2, log r2) they can have nonsensical long tails towards very low values. For visualization, we filtered out the long tail values (prior to selecting the 50 comparisons). The long tails tend to be most prevalent in the comparisons made from small feasible sets. Removing them therefore makes the small feasible sets look *more* self-similar.

#### R2

Higher R2 values indicate more similarity. R2 can be *very low* but the most meaningful variation is between 0 and 1.

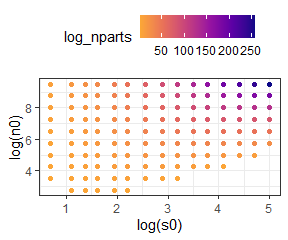

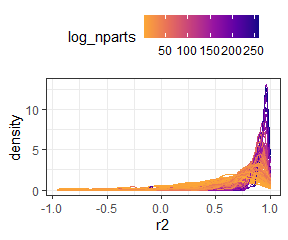

Here is R2 for only relatively small feasible sets (those with fewer than $e^{20}$ elements in the feasible set). This lets us zoom in on the distributions where they start to broaden out.

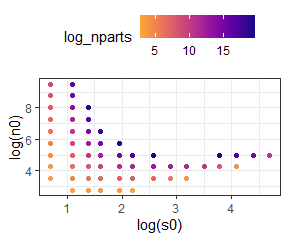

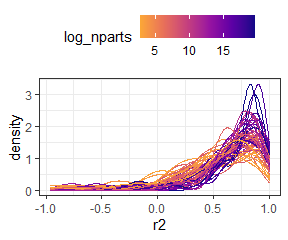

##### R2 on logged vectors

Higher R2 values indicate more similarity. R2 can be *very low* but the most meaningful variation is between 0 and 1.

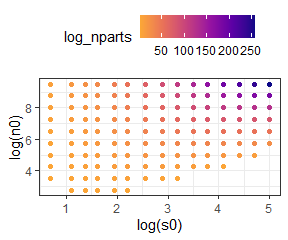

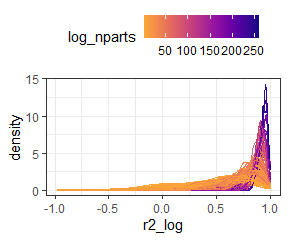

Here is R2 on logged vectors for only “small” feasible sets. This lets us zoom in on the distributions where they start to broaden out.

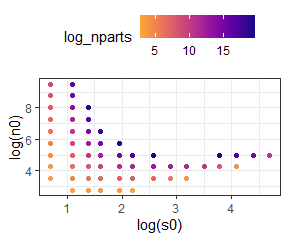

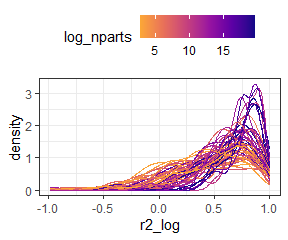

##### Coefficient of determination

Higher CD values indicate more similarity. It is bounded 0 to 1.

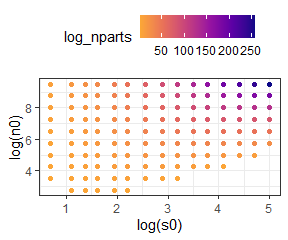

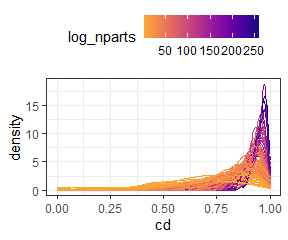

Here is cd for only “small” feasible sets.

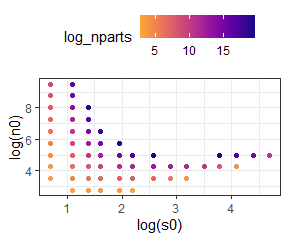

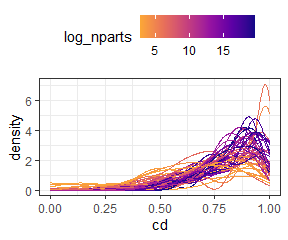

##### Proportion of individuals allocated to different species

Lower “prop off” values indicate more similarity. It is bounded 0 to 1.

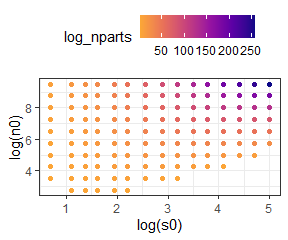

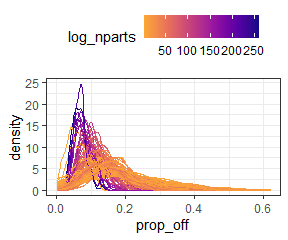

Here is prop_off for only “small” feasible sets.

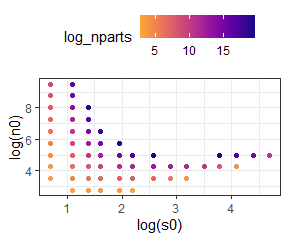

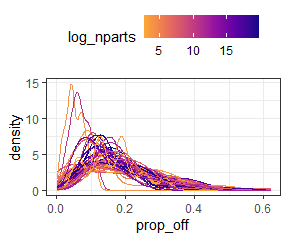

##### K-L divergence

Lower divergence values indicate more similarity. It is bounded 0 to 1.

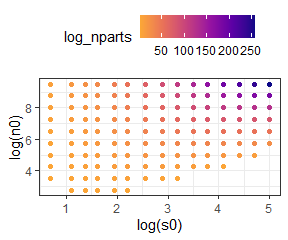

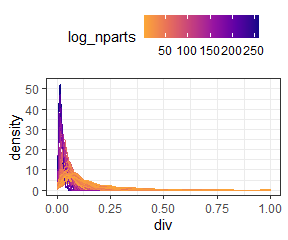

Here is K-L divergence for only “small” feasible sets.

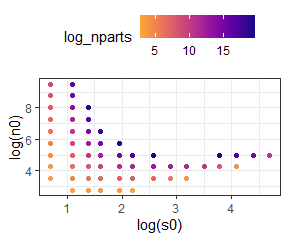

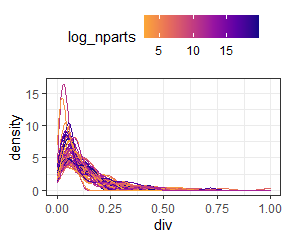

### In summary

Large feasible sets are consistently more self-similar than small ones, regardless of the metric of self-similarity used.
