## Appendix S7 for "Empirical abundance distributions are more uneven than expected given their statistical baseline"

Appendix A7: Complete results of resampling

2021-04-17

```
## Joining, by = c("sim", "source", "dat", "site", "singletons", "s0", "n0", "nparts")
## Joining, by = c("sim", "source", "dat", "site", "singletons", "s0", "n0", "nparts")
## Warning: Unknown or uninitialised column: `njks_skew`.
## `summarise()` has grouped output by 'Dataset'. You can override using the `.groups` argument.
## `summarise()` has grouped output by 'Dataset'. You can override using the `.groups` argument.
## Joining, by = c("Dataset", "resampling")
## Joining, by = "ometric2"
```

### Percentile histograms

#### Subsampling

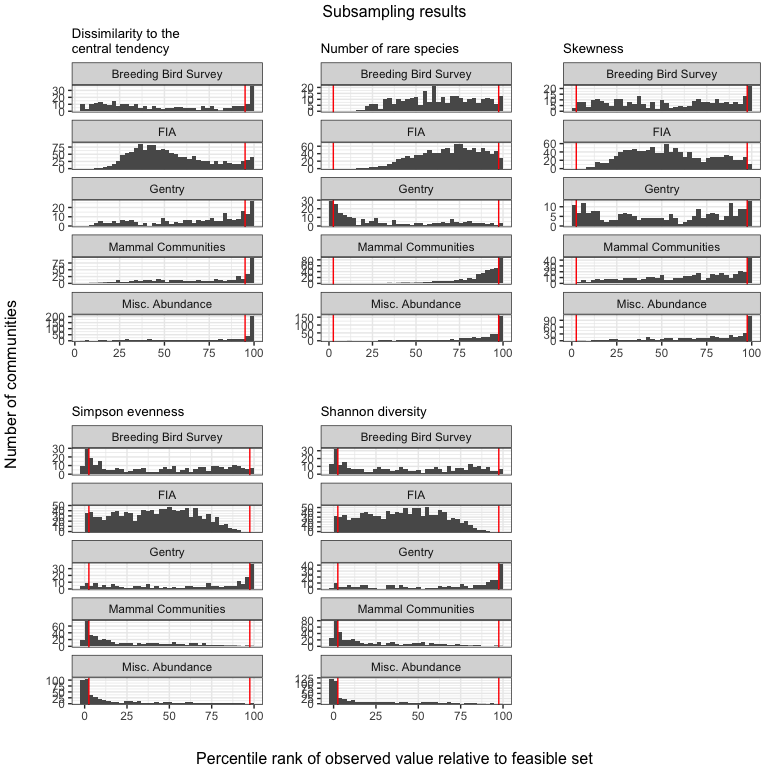

#### Adjusting for rare species

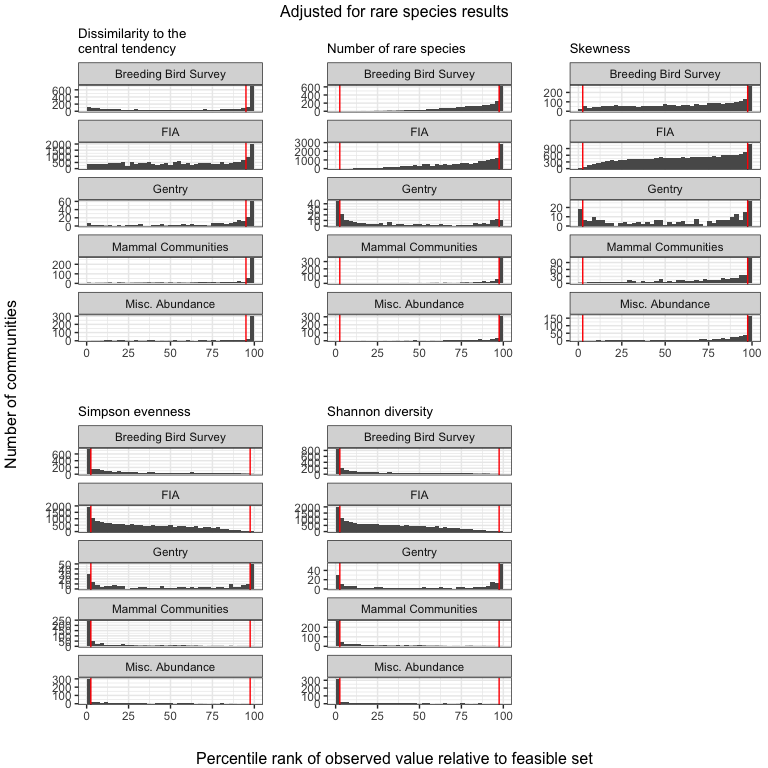

### Summary of effects on proportion of extreme values

#### Usual direction

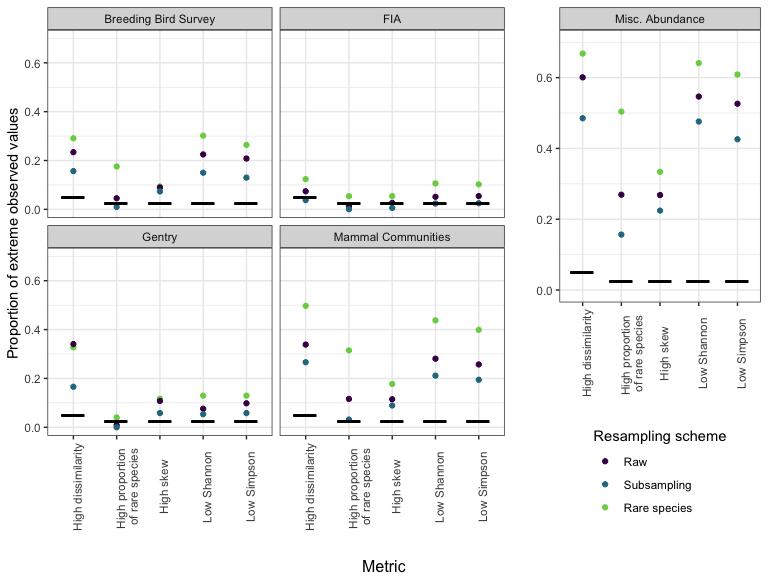

#### Unusual direction

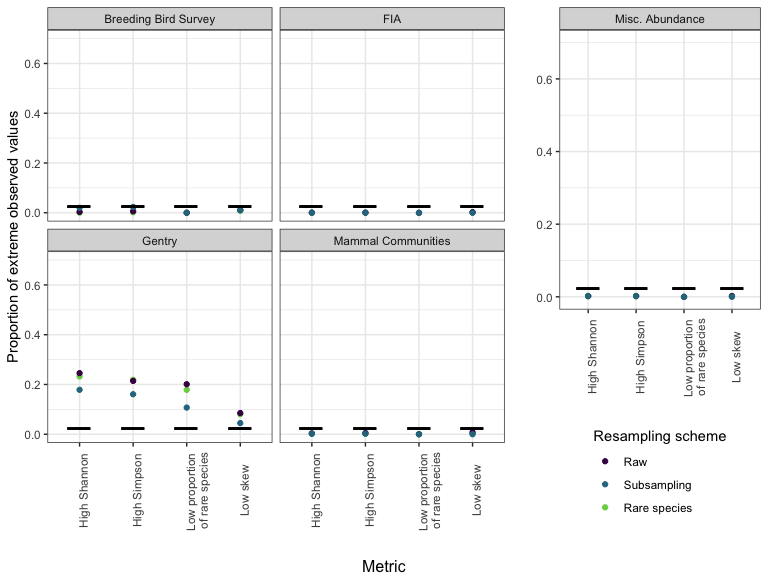

### Table of proportions of extreme values

#### Usual direction

```
## Note: Using an external vector in selections is ambiguous.
## ℹ Use `all_of(cols1)` instead of `cols1` to silence this message.
## ℹ See <https://tidyselect.r-lib.org/reference/faq-external-vector.html>.
## This message is displayed once per session.
```

| Dataset | Resampling scheme | High dissimilarity | High proportion of rare species | High skew | Low Simpson | Low Shannon |
| --- | --- | --- | --- | --- | --- | --- |
| Breeding Bird Survey | Raw | 23%; n =2773 | 4.5%; n =2773 | 9%; n = 2773 | 21%; n = 2773 | 23%; n = 2773 |
| Breeding Bird Survey | Subsampling | 16%; n =300 | 1%; n =300 | 7.3%; n = 300 | 13%; n = 300 | 15%; n = 300 |
| Breeding Bird Survey | Rare species | 29%; n =2773 | 18%; n =2773 | 9.4%; n = 2773 | 26%; n = 2773 | 30%; n = 2773 |
| FIA | Raw | 7.4%; n =17410 | 1.3%; n =18447 | 2.6%; n = 18447 | 5.4%; n = 18447 | 5.1%; n = 18447 |
| FIA | Subsampling | 3.8%; n =1118 | 0.072%; n =1388 | 0.58%; n = 1388 | 2.5%; n = 1388 | 2.4%; n = 1388 |
| FIA | Rare species | 12%; n =17918 | 5.4%; n =18736 | 5.4%; n = 18736 | 10%; n = 18736 | 11%; n = 18736 |
| Gentry | Raw | 34%; n =223 | 0.89%; n =224 | 11%; n = 223 | 9.8%; n = 224 | 7.6%; n = 224 |
| Gentry | Subsampling | 17%; n =223 | 0%; n =224 | 5.8%; n = 223 | 5.8%; n = 224 | 5.4%; n = 224 |
| Gentry | Rare species | 33%; n =223 | 4%; n =224 | 12%; n = 223 | 13%; n = 224 | 13%; n = 224 |
| Mammal Communities | Raw | 34%; n =511 | 12%; n =552 | 11%; n = 540 | 26%; n = 552 | 28%; n = 552 |
| Mammal Communities | Subsampling | 27%; n =432 | 3.2%; n =473 | 8.9%; n = 471 | 19%; n = 473 | 21%; n = 473 |
| Mammal Communities | Rare species | 50%; n =555 | 31%; n =594 | 18%; n = 592 | 40%; n = 594 | 44%; n = 594 |
| Misc. Abundance | Raw | 60%; n =486 | 27%; n =494 | 27%; n = 492 | 53%; n = 494 | 55%; n = 494 |
| Misc. Abundance | Subsampling | 49%; n =474 | 16%; n =479 | 22%; n = 477 | 43%; n = 479 | 48%; n = 479 |
| Misc. Abundance | Rare species | 67%; n =488 | 50%; n =496 | 33%; n = 494 | 61%; n = 496 | 64%; n = 496 |

#### Unusual direction

```
## Note: Using an external vector in selections is ambiguous.
## ℹ Use `all_of(cols2)` instead of `cols2` to silence this message.
## ℹ See <https://tidyselect.r-lib.org/reference/faq-external-vector.html>.
## This message is displayed once per session.
```

| Dataset | Resampling scheme | Low proportion of rare species | Low skew | High Simpson | High Shannon |
| --- | --- | --- | --- | --- | --- |
| Breeding Bird Survey | Raw | 0%; n =2773 | 1.1%; n = 2773 | 0.61%; n = 2773 | 0.36%; n = 2773 |
| Breeding Bird Survey | Subsampling | 0%; n =300 | 1%; n = 300 | 2.3%; n = 300 | 2%; n = 300 |
| Breeding Bird Survey | Rare species | 0%; n =2773 | 0.83%; n = 2773 | 0.22%; n = 2773 | 0.036%; n = 2773 |
| FIA | Raw | 0%; n =18447 | 0.27%; n = 18447 | 0.06%; n = 18447 | 0.081%; n = 18447 |
| FIA | Subsampling | 0%; n =1388 | 0%; n = 1388 | 0%; n = 1388 | 0%; n = 1388 |
| FIA | Rare species | 0%; n =18736 | 0.096%; n = 18736 | 0.011%; n = 18736 | 0.021%; n = 18736 |
| Gentry | Raw | 20%; n =224 | 8.5%; n = 223 | 21%; n = 224 | 25%; n = 224 |
| Gentry | Subsampling | 11%; n =224 | 4.5%; n = 223 | 16%; n = 224 | 18%; n = 224 |
| Gentry | Rare species | 18%; n =224 | 8.1%; n = 223 | 22%; n = 224 | 23%; n = 224 |
| Mammal Communities | Raw | 0%; n =552 | 0.74%; n = 540 | 0.54%; n = 552 | 0.36%; n = 552 |
| Mammal Communities | Subsampling | 0%; n =473 | 0%; n = 471 | 0.21%; n = 473 | 0.21%; n = 473 |
| Mammal Communities | Rare species | 0%; n =594 | 0.34%; n = 592 | 0.17%; n = 594 | 0.17%; n = 594 |
| Misc. Abundance | Raw | 0%; n =494 | 0.2%; n = 492 | 0.2%; n = 494 | 0.2%; n = 494 |
| Misc. Abundance | Subsampling | 0%; n =479 | 0%; n = 477 | 0.21%; n = 479 | 0.21%; n = 479 |
| Misc. Abundance | Rare species | 0%; n =496 | 0.2%; n = 494 | 0.2%; n = 496 | 0.2%; n = 496 |
