## Appendix S8 for "Empirical abundance distributions are more uneven than expected given their statistical baseline"

Unusual Gentry communities

Renata Diaz

2021-04-17

The Gentry communities are extreme relative to hte other datasets (in grey), in that they have high species richness and low abundance. Of these, the most extreme communities have extremely low average abundance (e.g. N/S < 3).

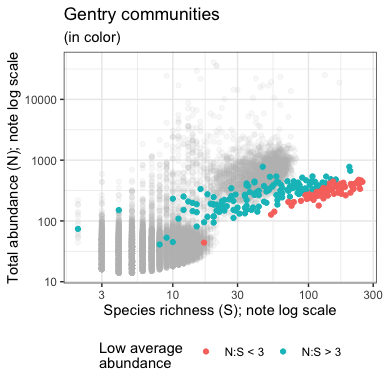

This figure shows all of the communities across all our datasets in terms of S and N. Gentry communities are in color, color coded by whether they have low average abundance (N/S < 3).

These low-average-abundance communities - the red dots - account for most of the signal of extreme values in the unusual direction for various metrics - that is, a **low** number of rare species and **low** skewness, and **high** Shannon and Simpson evenness. Gentry communities with average abundance > 3 tend to show the signal in the same direction as most other communities.

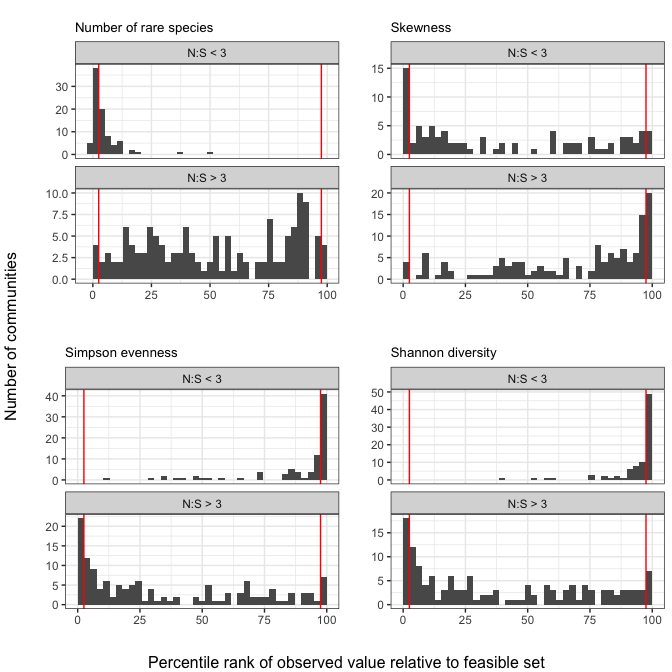
