## Appendix S9 for "Empirical abundance distributions are more uneven than expected given their statistical baseline"

Comparison of FIA and equally-sized communities

2021-04-17

We identified ~330 communities in FIA with exact matches, in terms of S and N, among communities from other datasets. We then compared the distributions of percentile scores and breadth indices of FIA communities to communities from other datasets, visually and using Kolmogorov-Smirnov tests.

### Percentile scores

#### Histograms

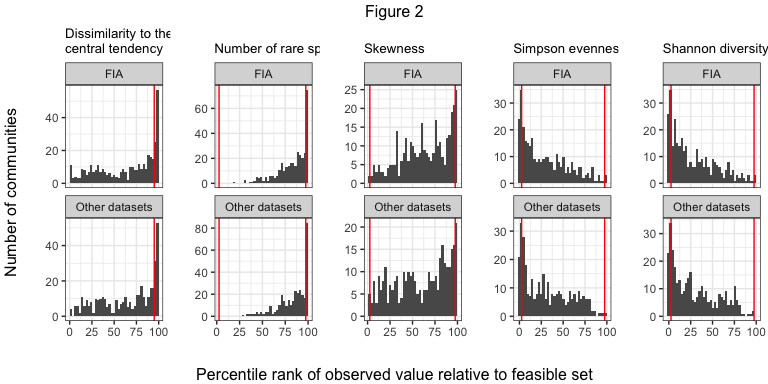

#### Proportions of extreme percentile scores

| Dataset | High dissimilarity | High proportion of rare species | High skew | Low Simpson | Low Shannon |
| --- | --- | --- | --- | --- | --- |
| FIA | 17%; n = 373 | 4.8%; n = 330 | 7%; n = 330 | 15%; n = 330 | 16%; n = 330 |
| Other datasets | 17%; n = 373 | 6.1%; n = 330 | 6.4%; n = 330 | 15%; n = 330 | 15%; n = 330 |

#### K-S test results

|  | Variable | K-S D | p-value |
| --- | --- | --- | --- |
| D…1 | Simpson | 0.0575758 | 0.6447379 |
| D…2 | Skew | 0.0000000 | 1.0000000 |
| D…3 | Shannon | 0.0606061 | 0.5794869 |
| D…4 | Number of rare species | 0.0363636 | 0.9812077 |
| D…5 | Dissimilarity to central tendency | 0.0424242 | 0.9277987 |

### Breadth indices

#### Histograms

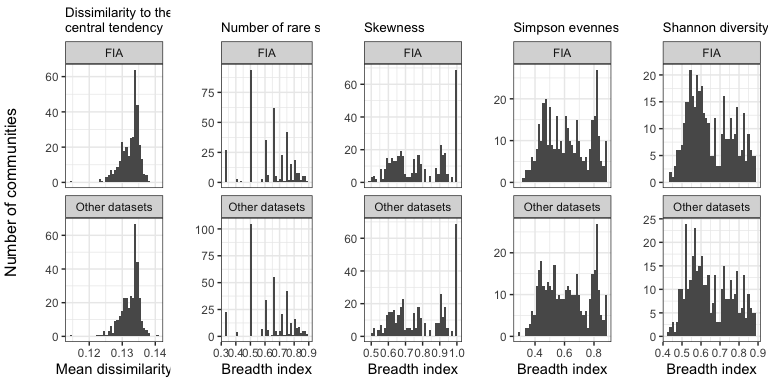

#### K-S test results

|  | Variable | K-S D | p-value |
| --- | --- | --- | --- |
| D…1 | Simpson | 0.0393939 | 0.9599607 |
| D…2 | Skew | 0.0000000 | 1.0000000 |
| D…3 | Shannon | 0.0424242 | 0.9277987 |
| D…4 | Number of rare species | 0.0333333 | 0.9930019 |
| D…5 | Dissimilarity to central tendency | 0.0268097 | 0.9993106 |
