## Appendix S10 for "Empirical abundance distributions are more uneven than expected given their statistical baseline"

Appendix A10: Complete results for very small communities

2021-04-17

We compare the results for very small communities (fewer than 2000 possible SADs in the feasible set) to larger communities within each dataset.

Only three of our datasets - Mammal Communities, Misc. Abundance, and FIA - have appreciable numbers of very small communities. We use 2000 possible SADs as a cutoff because it splits these datasets into reasonably large subsets of both “large” and “small” communities.

Proportions of communities in these datasets with fewer than 2000 possible SADs:

```
## `summarise()` has grouped output by 'Dataset'. You can override using the `.groups` argument.
## `summarise()` has grouped output by 'Dataset'. You can override using the `.groups` argument.
## `summarise()` has grouped output by 'Dataset'. You can override using the `.groups` argument.
## Joining, by = c("Dataset", "Number of elements")
## Joining, by = c("Dataset", "Number of elements")
```

| Dataset | Proportion with < 2000 possible SADs |
| --- | --- |
| FIA | 0.6804901 |
| Mammal Communities | 0.3496377 |
| Misc. Abundance | 0.0769231 |

### Proportions of communities with extreme percentile scores

The proportions of communities with >/< 2000 possible SADs with extreme percentile scores for each dataset.

For dissimilarity, approximately 5% of communities would have extreme scores by chance. For all other metrics, this would be approximately 2.5%.

In the direction of effects usually seen:

```
## Note: Using an external vector in selections is ambiguous.
## ℹ Use `all_of(cols1)` instead of `cols1` to silence this message.
## ℹ See <https://tidyselect.r-lib.org/reference/faq-external-vector.html>.
## This message is displayed once per session.
```

| Dataset | Number of elements | High dissimilarity | High proportion of rare species | High skew | Low Simpson | Low Shannon |
| --- | --- | --- | --- | --- | --- | --- |
| FIA | Less than 2000 | 5.3%; n = 12553 | 1.2%; n = 11516 | 1.2%; n = 11516 | 3.9%; n = 11516 | 3.6%; n = 11516 |
| FIA | More than 2000 | 11%; n = 5894 | 1.7%; n = 5894 | 6%; n = 5894 | 9.5%; n = 5894 | 9%; n = 5894 |
| Mammal Communities | Less than 2000 | 12%; n = 193 | 4.6%; n = 152 | 1.4%; n = 146 | 12%; n = 152 | 12%; n = 152 |
| Mammal Communities | More than 2000 | 42%; n = 359 | 16%; n = 359 | 17%; n = 359 | 35%; n = 359 | 38%; n = 359 |
| Misc. Abundance | Less than 2000 | 11%; n = 38 | 6.7%; n = 30 | 7.1%; n = 28 | 10%; n = 30 | 10%; n = 30 |
| Misc. Abundance | More than 2000 | 63%; n = 456 | 29%; n = 456 | 29%; n = 456 | 56%; n = 456 | 59%; n = 456 |

In the opposite direction:

```
## Note: Using an external vector in selections is ambiguous.
## ℹ Use `all_of(cols2)` instead of `cols2` to silence this message.
## ℹ See <https://tidyselect.r-lib.org/reference/faq-external-vector.html>.
## This message is displayed once per session.
```

| Dataset | Number of elements | Low proportion of rare species | Low skew | High Simpson | High Shannon |
| --- | --- | --- | --- | --- | --- |
| FIA | Less than 2000 | 0%; n = 11516 | 0.29%; n = 11516 | 0.069%; n = 11516 | 0.096%; n = 11516 |
| FIA | More than 2000 | 0%; n = 5894 | 0.27%; n = 5894 | 0.051%; n = 5894 | 0.068%; n = 5894 |
| Mammal Communities | Less than 2000 | 0%; n = 152 | 1.4%; n = 146 | 0%; n = 152 | 0%; n = 152 |
| Mammal Communities | More than 2000 | 0%; n = 359 | 0.56%; n = 359 | 0.84%; n = 359 | 0.56%; n = 359 |
| Misc. Abundance | Less than 2000 | 0%; n = 30 | 0%; n = 28 | 0%; n = 30 | 0%; n = 30 |
| Misc. Abundance | More than 2000 | 0%; n = 456 | 0.22%; n = 456 | 0.22%; n = 456 | 0.22%; n = 456 |

### Plots of breadth indices and percentile scores

Breadth indices (top) and percentile scores (bottom) for communities with fewer than 2000 possible SADs, or more than 2000 possible SADs, from FIA, Mammal Community, and Misc. Abundance datasets.

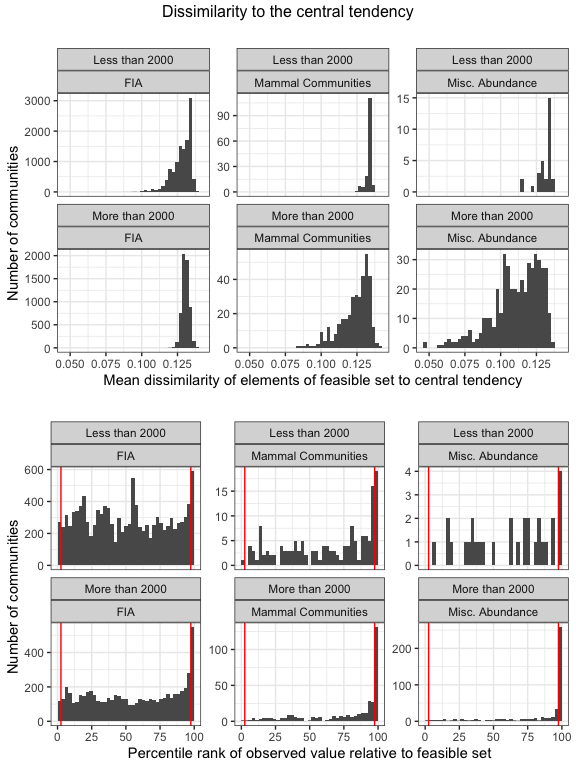

\## Warning: Removed 14 rows containing non-finite values (stat_bin).

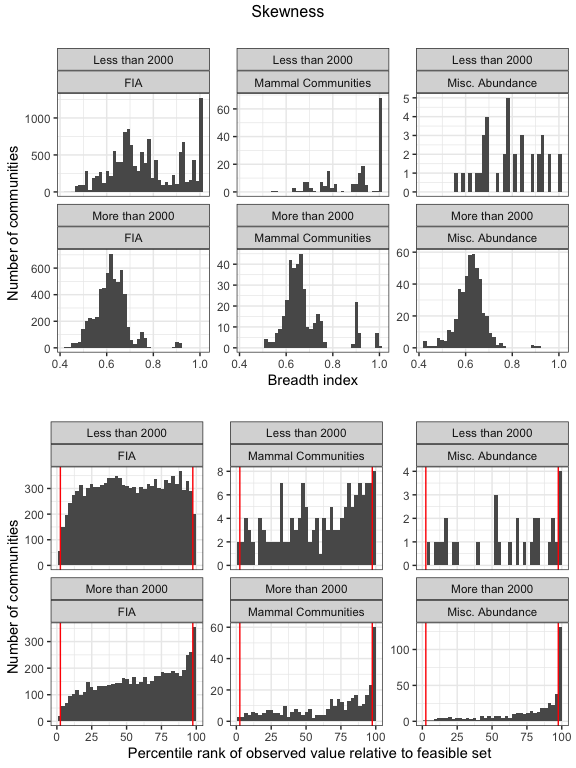

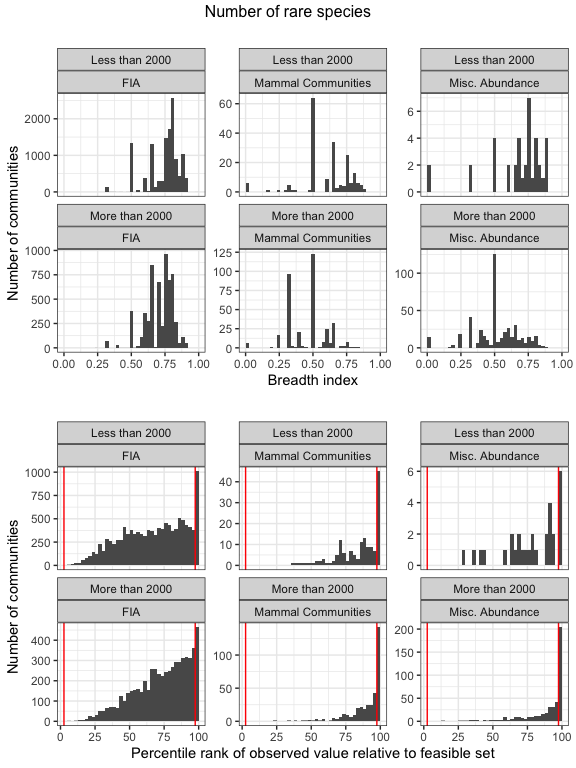

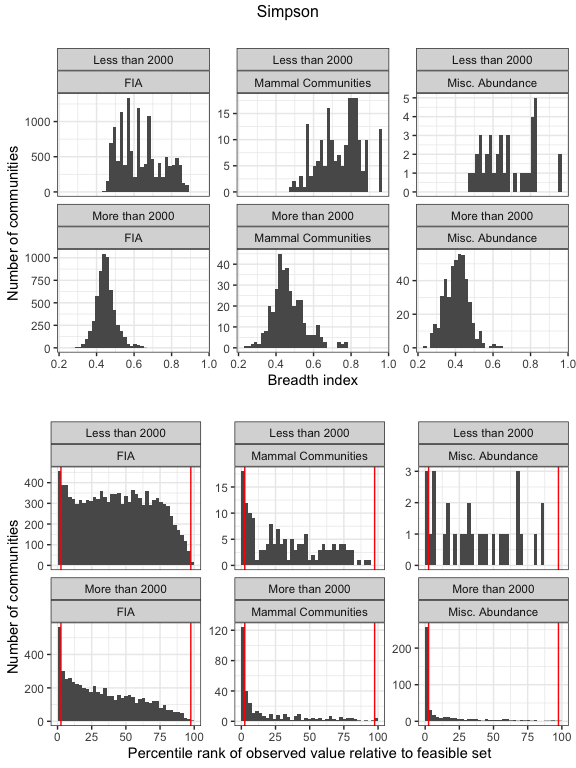

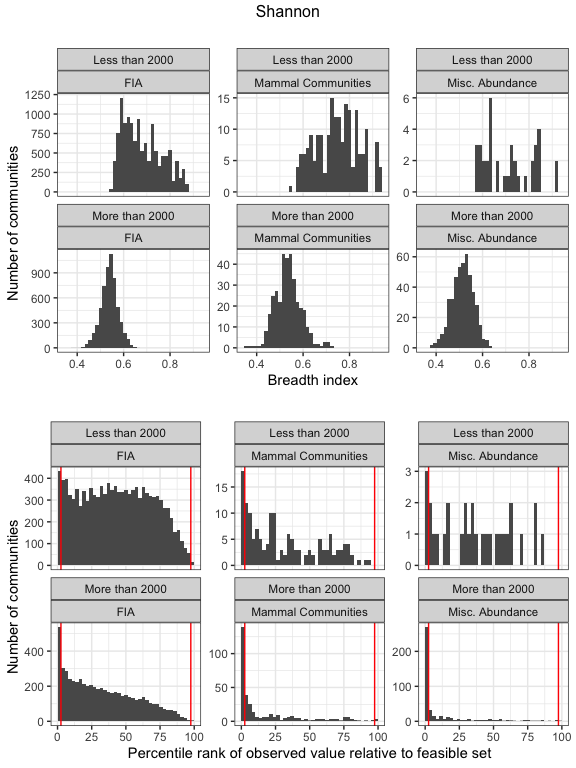
