## Supplementary material for "Empirical abundance distributions are more uneven than expected given their statistical baseline": Figure S4

2021-04-17

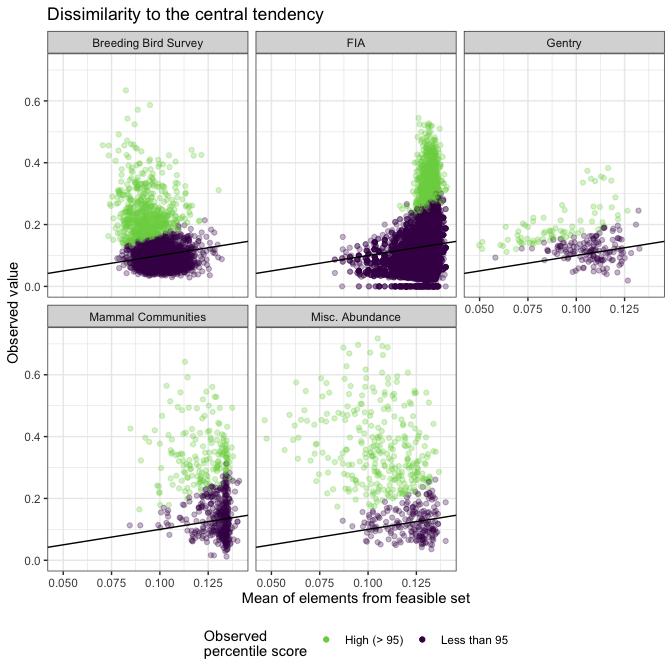

**Figure S4**. Dissimilarity of observed and sampled SADs to the central tendency of the feasible set. Observed SADs are often much more dissimilar to the central tendency of their feasible sets (y-axis) than the mean dissimilarity of samples from the feasible set and the central tendency (x-axis). Dissimilarity can range from 0-1. The black line is the 1:1 line. Of observed SADs that are more disssimilar to the central tendency than are 95% of samples from the feasible set (green points), the absolute increase in dissimlarity ranges from .04 to .6. These observed SADs are from 1.4 to 9.7 times more dissimilar to the central tendency than the maen of samples from the feasible set.
