## Supplementary material for "Empirical abundance distributions are more uneven than expected given their statistical baseline": Figure S6

2021-04-17

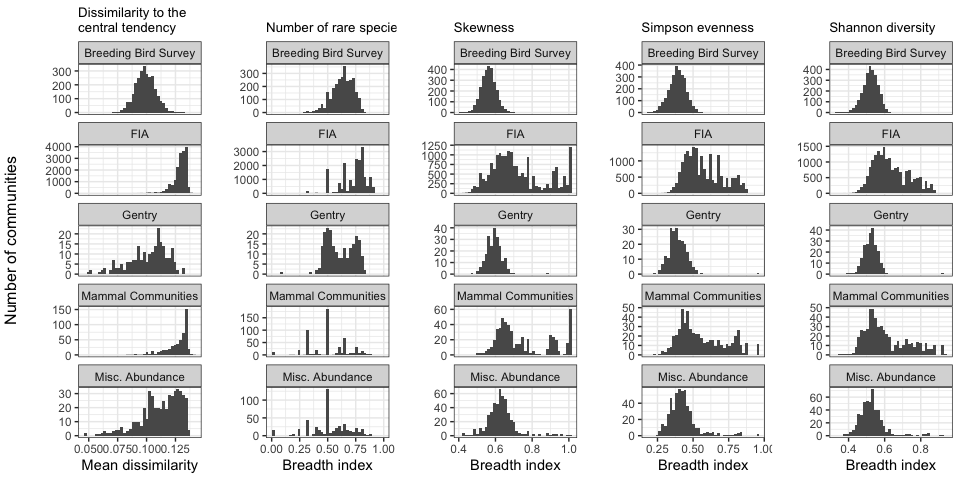

**Figure S6**. Partially because of the uneven distribution of S and N among the different datasets, the narrowness of the feasible sets - defined either as the mean dissimilarity of samples from the feasible set to the central tendency of the feasible set, or using a breadth index for specific metrics - varies among different datasets. In particular, the FIA dataset, and subsets of the Mammal Community and Miscellaneous Abundance databases, often have highly variable, broadly-defined statistical baselines derived from the feasible set.
