## Supplementary material for "Empirical abundance distributions are more uneven than expected given their statistical baseline": Table S5

Proportions of extreme values the unusual direction

2021-04-17

```
## Joining, by = "Dataset"
## Joining, by = "Dataset"
## Note: Using an external vector in selections is ambiguous.
## ℹ Use `all_of(cols2)` instead of `cols2` to silence this message.
## ℹ See <https://tidyselect.r-lib.org/reference/faq-external-vector.html>.
## This message is displayed once per session.
```

| Dataset | Low proportion of rare species | Low skew | High Simpson | High Shannon |
| --- | --- | --- | --- | --- |
| Breeding Bird Survey | 0%; n = 2773 | 1.1%; n = 2773 | 0.61%; n = 2773 | 0.36%; n = 2773 |
| FIA | 0%; n = 17410 | 0.28%; n = 17410 | 0.063%; n = 17410 | 0.086%; n = 17410 |
| Gentry | 20%; n = 223 | 8.5%; n = 223 | 22%; n = 223 | 25%; n = 223 |
| Mammal Communities | 0%; n = 511 | 0.79%; n = 505 | 0.59%; n = 511 | 0.39%; n = 511 |
| Misc. Abundance | 0%; n = 486 | 0.21%; n = 484 | 0.21%; n = 486 | 0.21%; n = 486 |

**Table S5.** Proportions of extreme values for percentile scores for observed SADs compared to samples from the feasible set, for the less-usual direction of effects. This is the proportion of scores <2.5 or >97.5; by chance ~2.5% of scores should be in either extreme. n refers to the number of communities included for each dataset for each metric.
